# Screening the human druggable genome identifies ABHD17B as an anti-fibrotic target in hepatic stellate cells

**DOI:** 10.1101/2023.08.07.551744

**Authors:** Wenyang Li, Robert P. Sparks, Cheng Sun, Yang Yang, Lorena Pantano, Rory Kirchner, Arden Weilheimer, Benjamin J. Toles, Jennifer Y. Chen, Sean P. Moran, Victor Barrera, Zixiu Li, Peng Zhou, Meghan L. Brassil, David Wrobel, Shannan J. Ho Sui, Gary Aspnes, Michael Schuler, Jennifer Smith, Benjamin D. Medoff, Chan Zhou, Carine M. Boustany-Kari, Jörg F. Rippmann, Daniela M. Santos, Julia F. Doerner, Alan C. Mullen

## Abstract

Hepatic stellate cells (HSCs) are activated with chronic liver injury and transdifferentiate into myofibroblasts, which produce excessive extracellular matrices that form the fibrotic scar. While the progression of fibrosis is understood to be the cause of end-stage liver disease, there are no approved therapies directed at interfering with the activity of HSC myofibroblasts. We performed a high-throughput small interfering RNA (siRNA) screen in primary human HSC myofibroblasts to identify gene products necessary for the fibrotic phenotype of HSCs. We found that depletion of *ABHD17B* promotes the inactivation of HSCs, characterized by reduced *COL1A1* and *ACTA2* expression and accumulation of lipid droplets. Mice deficient for *Abhd17b* are also protected from fibrosis in the setting of *in vivo* liver injury. While ABHD17B is a depalmitoylase, our data suggest that ABHD17B promotes fibrosis through pathways independent of depalmitoylation that include interaction with MYO1B to modulate gene expression and HSC migration. Together, our results provide an analysis of the phenotypic consequences for siRNAs targeting RNAs from >9,500 genes in primary human HSCs and identify ABHD17B as a potential therapeutic target to inhibit liver fibrosis.

## Introduction

End-stage liver disease results from progressive fibrosis in the setting of chronic liver injury. While reversal of the underlying cause of injury can reduce the severity of fibrosis^1^, there are limited treatments for many sources of chronic injury, including non-alcoholic fatty liver disease/metabolic dysfunction-associated steatotic liver disease (NAFLD/MASLD)^2^. In other diseases of fibrotic injury, such as primary sclerosing cholangitis (PSC), there are no effective treatments beyond liver transplantation^3^.

HSC myofibroblasts are the primary cell type responsible for liver fibrosis^4–6^. In a healthy liver, HSCs are in a quiescent state and store vitamin A in lipid droplets^4^. With chronic injury, HSCs are activated and transdifferentiate into HSC myofibroblasts, characterized by the loss of lipid droplets and production of extracellular matrix (ECM) proteins that form the fibrotic scar^7,8^.

Resolution of liver fibrosis has been observed when the source of liver injury is removed^9^. In this setting, the HSC population is reduced through apoptosis, while 40-50% of HSC myofibroblasts revert to an inactive state, characterized by decreased collagen expression^10,11^. This suggests that with the correct signals, HSC myofibroblasts can be directed to an inactive phenotype.

Here, we performed an siRNA screen of the human druggable genome and >2,000 long noncoding (lnc) RNAs to identify RNAs that could be targeted to promote HSC inactivation. We found that depletion of *ABHD17B* (*FAM108B1*) in primary human HSCs led to a reduction in type I collagen expression and promoted HSC inactivation. These functions were independent of the depalmitoylase activity of ABHD17B and involve interaction with MYO1B. Furthermore, mice deficient in *Abhd17b* were protected from the development of liver fibrosis.

## Results

### Development of a high-throughput siRNA screen in primary human HSC myofibroblasts

We first optimized siRNA transfection conditions for human HSC myofibroblasts in a 384-well format (**Supplementary Fig. 1a, b**) by adapting approaches previously established for small molecule screens^12,13^. Next, we evaluated negative and positive controls. We transfected HSCs with non-targeting control siRNAs (NTC si) 1, 2, 3, 4 and 5 and performed Bodipy and Hoechst staining to evaluate the effect of the siRNAs on lipid accumulation and cell number. Compared to NTC si1 and si2, HSCs transfected with NTC si3, si4, and si5 demonstrated lower percentages of Bodipy positive cells with similar or reduced effect on cell numbers (**Supplementary Fig. 1c, d**). We compared NTC si3-si5 siRNAs with pooled siRNAs targeting *GAPDH* (*GAPDH* siP). Transfection with *GAPDH* siP did not cause a significant change in lipid accumulation (**Supplementary Fig. 1e**) and did not reduce cell numbers to the degree observed with NTC siRNAs (**Supplementary Fig. 1f**). Based on these results, we selected *GAPDH* siP as our negative control.

We then evaluated potential positive controls for induction of lipid droplets and transfection efficiency. siRNAs targeting *ACTA2* and *ASAH1*^12^ did not significantly increase the percentage of Bodipy-positive cells (**Supplementary Fig. 1c**). Depletion of *PLK1* via siRNAs is established to cause death in multiple cell lines^14^, but despite ∼70% depletion, reduction in HSC number was mild (∼30%-40% decrease) (**Supplementary Fig. 1g, h**). These results showed that depletion of *ACTA2* or *ASAH1* did not affect lipid accumulation, and depletion of *PLK1* did not affect cell number sufficiently for these siRNAs to be used as controls.

To identify additional siRNA controls, we screened siRNA SMARTpools defined as hits in at least two different siRNA screens at the Institute of Chemistry and Cell Biology-Longwood (ICCB-L) (**Supplementary Fig. 2a**). We calculated the Z’ factor for each plate based on the percent positive cells of nortriptyline^12^ and *GAPDH* SMARTpool control wells and confirmed that the Z’ was acceptable (0.4-0.5). We identified potential positive hits, including *AURKC*, *AKAP11*, and *TLR6* siRNAs (**Supplementary Fig. 2a**). We tested these siRNAs, and siRNAs targeting *NFkB1*, *NFkB2*, *NR1H2,* and *NR1H3*, which are annotated target genes shared by positive hits from our small molecule screen^13^, for potential positive controls. Unfortunately, none of these siRNAs showed a sufficiently robust phenotype to serve as a positive control for Bodipy staining (**Supplementary Fig. 2b-j**). Depletion of *UBB* did dramatically reduced cell number (**Supplementary Fig. 2a, Supplementary Fig. 3a, b**). We tested the Z’ of *UBB* siP versus *GAPDH* siP and nortriptyline versus *GAPDH* siP and confirmed that the Z’ values (0.25 and 0.60) were acceptable (**Supplementary Fig. 3c, d**). *UBB* pooled siRNA was selected as a control for transfection efficiency and nortriptyline^12,13^ as a positive control for Bodipy staining.

### Primary siRNA screen identifies pooled siRNAs that inactivate HSC myofibroblasts

We screened the Dharmacon SMARTpool siRNA library targeting a total of 7641 mRNAs from the human druggable genome as well as siRNAs targeting 2237 lncRNAs in a one target/well format. Each SMARTpool was composed equally of four duplexes targeting the same RNA. The screen was conducted in technical triplicate (three distinct wells for each siRNA pool), with *GAPDH* SMARTpool siRNA as the negative control, *UBB* SMARTpool siRNA as the control for transfection efficiency, and 15 μM nortriptyline as the control for lipid accumulation and Bodipy staining (**Fig. 1a, b**). As indicated by the high toxicity of *UBB* siRNAs and high score of nortriptyline, the transfection was efficient and the staining worked as expected **(Fig. 1a**). The screening results for all siRNAs and controls are provided in **Supplementary Table 1**.

**Fig. 1.**
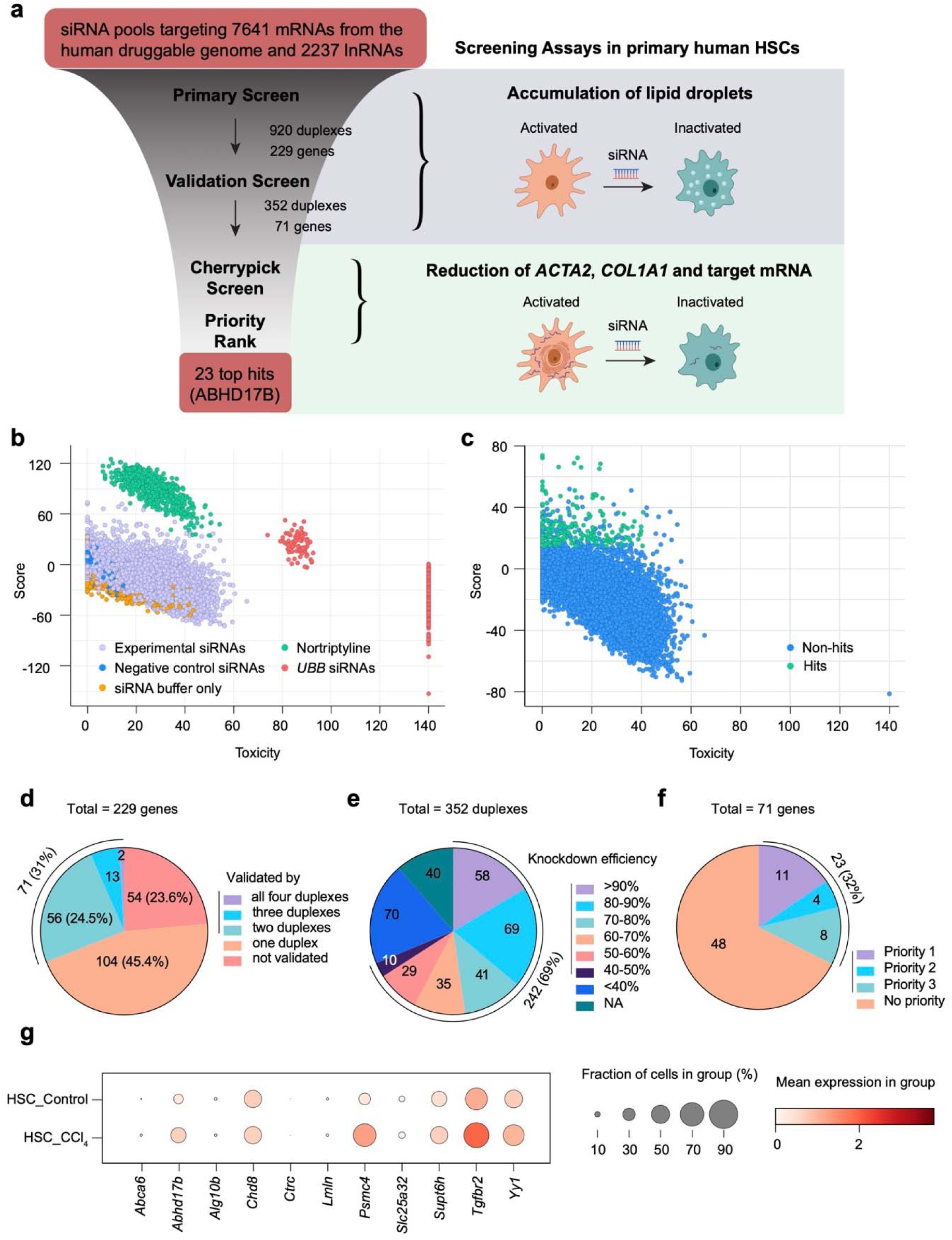
High-throughput screening identified siRNAs that induce HSC myofibroblast inactivation. (a) Overview of the siRNA screen. (b) Results of the primary siRNA screen. Each dot represents the integrated score and toxicity value of three replicates of an experimental or control well. Data from all wells are shown. (c) The score and toxicity values for all experimental wells from the primary screen. (d) Summary of the validation screen results. (e) Summary of the siRNA knockdown efficiency in the cherry pick screen. (f) Grouping the final 71 hits by priority. (g) Expression of the 11 priority 1 genes in HSCs from the livers of CCl_4_-treated and control mice based on single-cell RNA sequencing results^16^. Circle size represents the fraction of cells expressing a gene, and color indicates mean expression level.

We adapted our previous analysis method to calculate the score and toxicity for each siRNA pool ^13^, based on the average percent Bodipy-positive cells for each siRNA pool compared to the baseline of the plate, the correlation among the three replicates, and total cell numbers. There was a small fraction of outliers, possibly due to inefficient depletion for a replicate or artifacts acquired during image acquisition. We developed a computational algorithm to correct outliers **(Supplementary Methods)**, which allowed us to retain variability while reducing the extreme variance that an outlying replicate could generate.

We identified 229 unique gene products as primary hits. This corresponded to 231 positive experimental wells because two wells contained the same siRNA pool, and one gene product was targeted by two different siRNAs. We selected primary hits based on the following criteria: <u>1</u>. the score was significantly increased (FDR < 0.05); <u>2</u>. toxicity was lower than the threshold determined by the nortriptyline wells; and <u>3a</u>. transcripts per million (TPM) > 1 (for mRNAs) or TPM > 0.5 (for lncRNAs) in at least one of 18 HSC RNA sequencing samples^12,15^ or <u>3b</u>. no sequencing data were available in these analyzed datasets (**Fig. 1c**). The screening results and expression analysis for all selected genes are provided in **Supplementary Table 2**.

### Secondary screen with individual siRNAs

We next examined the phenotypic consequences of target depletion with individual duplexes in a one duplex/well, four wells/target format. This deconvolution secondary screen consisted of 920 experimental duplexes (targeting transcripts from 229 genes) tested in technical triplicate, relying on the same assay, controls, and analysis strategy as the primary screen (**Supplementary Table 3**). A total of 71 transcripts/gene products were validated as defined by at least two duplexes increasing Bodipy staining with an FDR < 0.05 and toxicity lower than the threshold determined by the *UBB* depletion (**Fig. 1d, and Supplementary Table 4**).

### Screening for hits that regulate ACTA2 and COL1A1

We next sought to quantify reduction in *ACTA2* and *COL1A1* as markers of HSC inactivation for the 71 genes identified in the deconvolution screen. For each condition we also quantified depletion of the mRNA targeted by each siRNA. To minimize technical variations, we measured *ACTA2*/*COL1A1*/target mRNA level in the same reaction as the housekeeping control *PSMB2*^13^ by performing multiplexed qRT-PCR with probes labeled with different dyes. In addition to the two duplexes with the highest scores in the deconvolution screen, we included three additional siRNA duplexes from the Ambion siRNA library for 70 genes (one of the genes was not in the Ambion library). 352 siRNA duplexes were tested by qRT-PCR in quadruplicate.

We analyzed mRNA levels for *ACTA2 and COL1A1* using two methods: a linear regression method and the standard ΔΔCt method with *PSMB2* as the endogenous control (**Supplementary Table 5, Supplementary Methods**). *COL1A1* reduction did not always correlate with *ACTA2* reduction. Approximately 66% of the siRNA duplexes showed a knockdown efficiency >50%, and ∼69% of the siRNA duplexes reduced the expression of the target transcript by more than 40%. The knockdown efficiency for 11% of siRNA duplexes could not be determined due to low expression of the target transcript or low-quality melting curves (**Fig. 1e**). We prioritized candidates based on the following criteria: **priority 1**: at least two siRNAs reduced *COL1A1* by at least 30% (based on the fold change calculated by either method) with at least 40% depletion of the target gene; **priority 2**: at least three siRNAs reduced *ACTA2* by at least 60% (fold change calculated by either method) with at least 40% depletion of the target gene and candidate was not included in priority 1; **priority 3**: two siRNAs reduced *ACTA2* by at least 60% (fold change calculated by either method) with at least 40% depletion of the target gene and candidate was not included in priority 1 or 2. We identified eleven genes as priority 1, four as priority 2, and eight as priority 3 (**Fig. 1f** and **Table 1**).

**Table 1.**
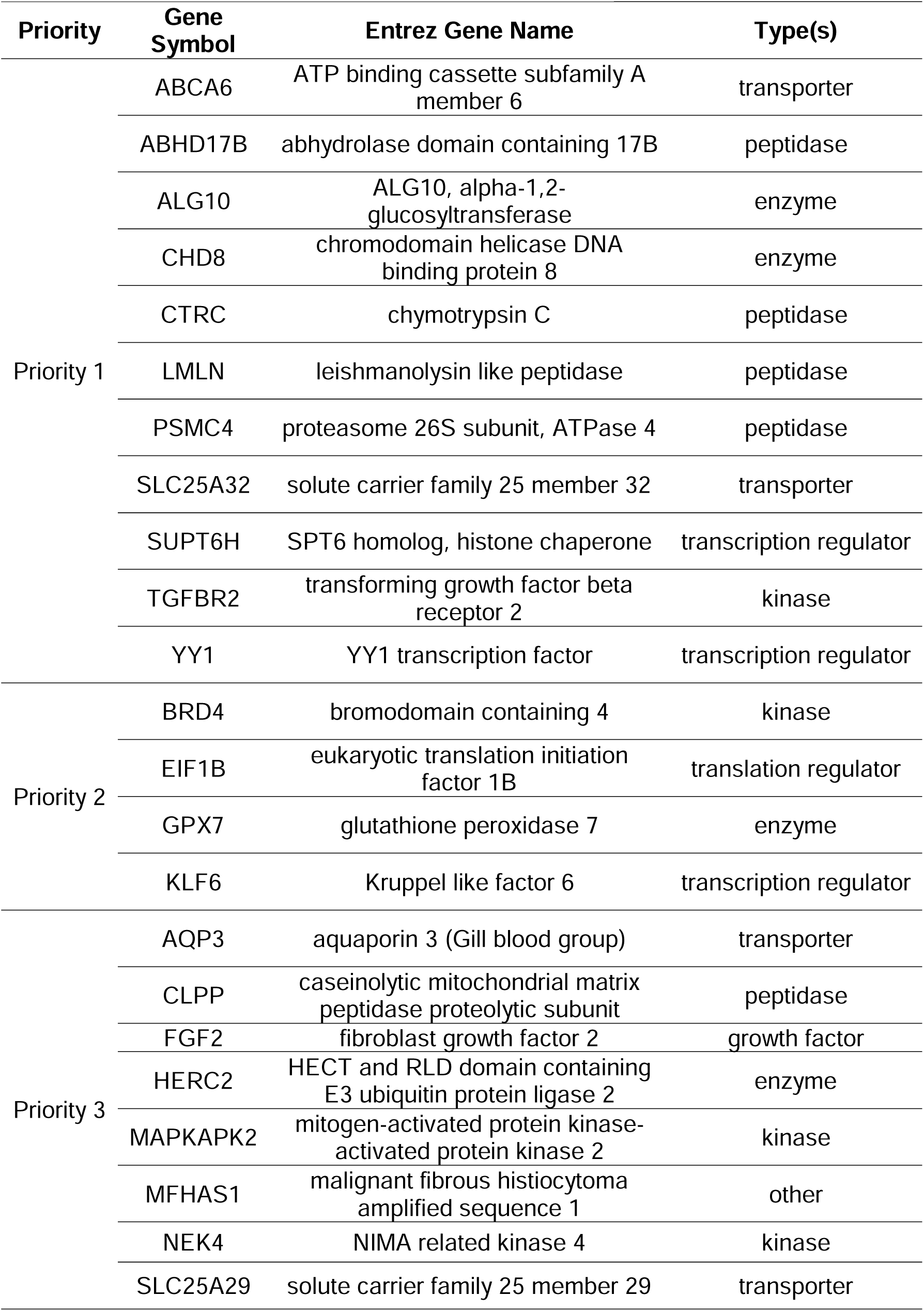
siRNA candidates grouped by priority.

Of note, 37 lncRNAs were among the 229 hits in the primary screen, and three lncRNAs (*AK2*/NR_037592, *C1orf140*/NR_024236, and *CLTB*/NR_045724) were among the 71 hits in the deconvolution screen. Depletion of these lncRNA genes had some effect on *ACTA2* but did not reduce *COL1A1* mRNA levels, and no lncRNA genes were ranked as priority 1-3 candidates.

### ABHD17B depletion increases lipid accumulation and induces HSC inactivation

We further analyzed the expression of the priority 1 genes in fibrotic livers based on single-cell RNA sequencing (scRNA-seq) data from control and CCl_4_-treated mice^16^. Only *Abhd17b, Tgfbr2,* and *Psmc4* were expressed in more than 10% of HSCs and demonstrated an average increase in expression of greater than 1.5 fold in HSCs with CCl_4_ treatment **(Fig. 1g and Supplementary Table 6)**. Interfering with Tgfbr2 activity or blocking TGF-β signaling in HSCs reduces liver fibrosis *in viv*o^17,18^, supporting inclusion of *TGFBR2* in the final group of genes. We also found that depletion of *PSMC4,* a subunit of the 26S proteasome^19^, reduced *ACTA2* and *COL1A1* expression in primary human HSCs from a second donor (**Supplementary Fig. 4**), further confirming results from the screen.

*ABHD17B* encodes an enzyme with palmitoyl-hydrolase activity, and members of the ABHD17 family regulate palmitoylation of proteins including N-Ras^20^. The function of ABHD17B in HSCs in unknown, but in models of chronic injury, it is induced to a higher degree in HSCs than hepatocytes or any other nonparenchymal cell type with the exception of lymphocytes (**Supplementary Fig. 5)**. We confirmed the effect of *ABHD17B* depletion on lipid accumulation using primary HSCs from multiple donors. Transfection with two different *ABHD17B* siRNA duplexes significantly increased the accumulation of lipid droplets in cells from three donors. HSCs from the remaining donor (2) showed a significant increase in lipid accumulation with one *ABDH17B* siRNA (**Fig. 2a**). These results confirm that depletion of *ABHD17B* is sufficient to increase lipid accumulation, which suggests an inactive HSC phenotype.

**Fig. 2.**
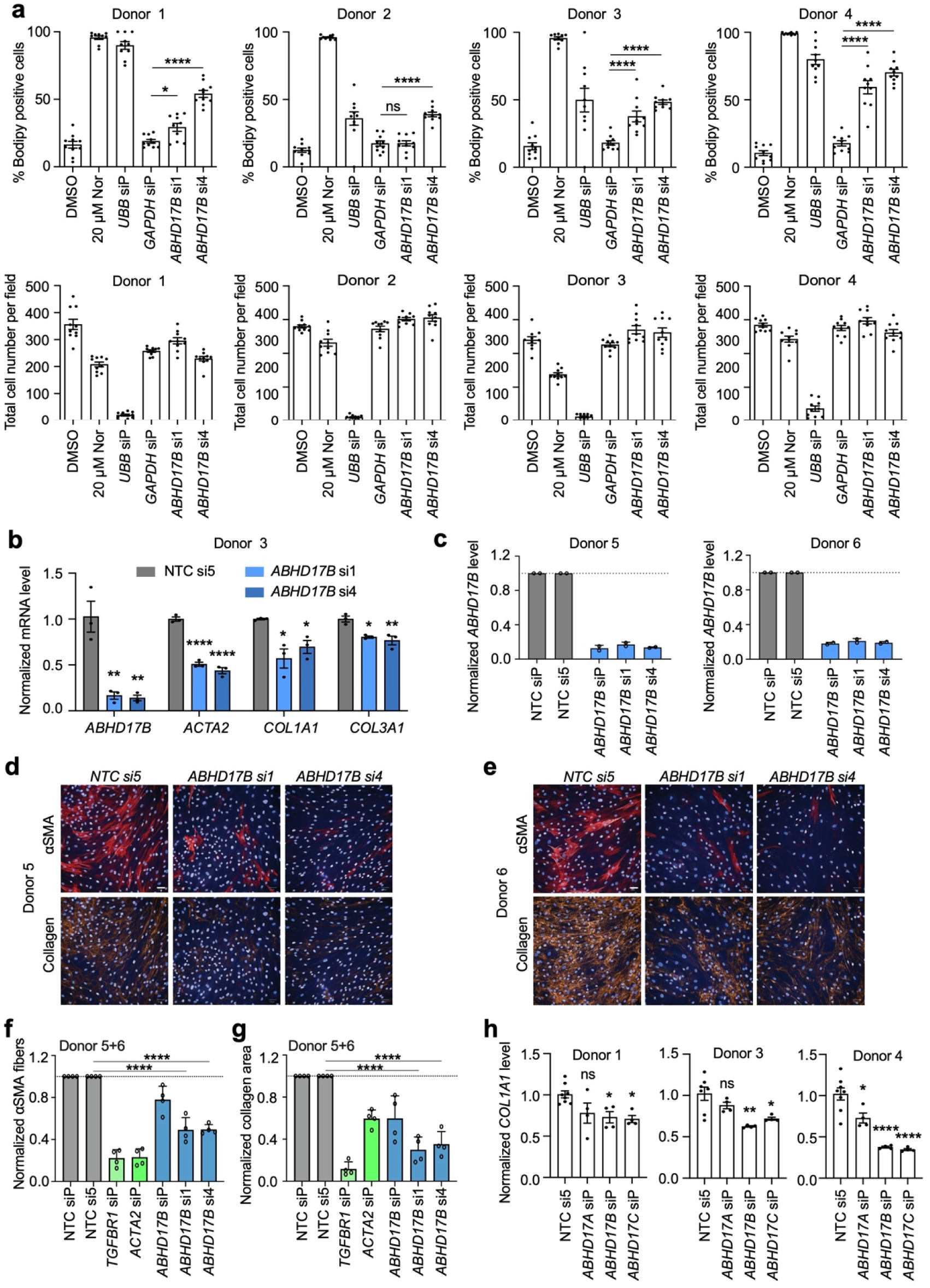
Depletion of *ABHD17B* inactivates HSCs. (a) The fraction of Bodipy positive cells (top) and total cell numbers (bottom) are showed for HSCs transfected with siRNAs targeting *ABDH17B*. DMSO and nortriptyline (Nor) serve as negative and positive controls for Bodipy, respectively. siRNA targeting *UBB* serve as a control for transfection efficiency and siRNAs targeting *GAPDH* as an siRNA negative control for Bodipy. Each dot represents one biological replicate. Error bars represent mean ± SEM (n=10). ns indicates not significant, * indicates p<0.05, and **** indicates p<0.0001 (one-way ANOVA test). (b) *ACTA2*, *COL1A1,* and *COL3A1* expression was quantified by qRT-PCR two days after HSCs were transfected with the indicated siRNAs. Each dot represents one biological replicate. Error bars represent mean ± SEM (n=3). * indicates p<0.05, ** indicates p<0.01, and **** indicates p<0.0001 (one-way ANOVA test). Data are representative of three independent experiments. (c) *ABDH17B* expression was quantified by qRT-PCR three days after transfection in HSCs (n=2). Non-targeting control pooled siRNA (NTC siP) and NTC si5 are compared to pooled ABHD17B (siP) siRNAs, *ABHD17B si1*, and *ABHD17B si4*. (d-e) HSCs were transfected with the indicated siRNAs prior to serum starvation and stimulation with scar-in-a-jar conditions for 72 hr. α−SMA (red) and collagen (orange) were visualized by immunofluorescence in the same field of view. Nuclei were stained with Hoechst (blue). White bars indicate 50 µm. (f-g) Quantification of α-SMA and collagen from scar-in-a-jar assay. Normalized α-SMA fiber score is displayed for indicated siRNAs in (f). Normalized quantification of collagen area is shown in (g). Data from donors 5 and 6 are combined. Error bars represent mean ± SD (n=4). **** indicates p<0.0001. p<0.0001 for *TGFBR1 siP* and *siACTA2 siRNA* compared *to pooled NTC siP,* and p <0.01 for *ABHD17B siP* compared to *NTC siP* (Dunnetts multiple comparison test). (h) Members of the ABHD17 family were depleted using pooled siRNAs. *COL1A1* levels were quantified by qRT-PCR three days after transfection. Each dot represents one biological replicate. Error bars represent mean ± SEM (n≥4). ns indicates not significant, * indicates p<0.05, ** indicates p<0.01, and **** indicates p<0.0001 (one-way ANOVA test).

Consistent with screening results in HSCs (**Fig. 1**), depletion of *ABHD17B* reduced *ACTA2*, *COL1A1,* and *COL3A1* levels in HSCs from a second donor (**Fig. 2b**). Next, we tested if depletion of *ABHD17B* affects the formation of α-SMA fibers as a marker of HSC activation and collagen deposition in the ECM as an indication of fibrotic activity. Both the siRNA pool (siP) and two different siRNA duplexes targeting *ABHD17B* significantly reduced *ABHD17B* mRNA in HSCs from two different donors (**Fig. 2c**) without affecting cell number (**Supplementary Fig. 6a**), resulting in HSC inactivation as evident from reduced staining of α-SMA (encoded by *ACTA2*) and collagen type I (**Fig. 2d-g, Supplementary Fig. 6b, c**). siRNAs targeting *TGFBR1* and *ACTA2* were included as positive controls. The observed effect of *ABHD17B* depletion on HSC phenotype suggests that ABHD17B promotes maintenance of myofibroblast features, and depletion leads to inactivation.

### ABHD17B promotes collagen expression in lung fibroblasts

Similar to liver fibrosis, idiopathic pulmonary fibrosis (IPF) is characterized by excessive ECM synthesized by activated fibroblasts and has limited treatment options^21^. We asked if ABHD17B also plays a role in promoting the fibrotic activity of primary human lung fibroblasts. We found that depletion of *ABHD17B* significantly reduced the expression of *COL1A1* and *ACTA2* (**Supplementary Fig. 7**), indicating that ABHD17B also plays a role in regulating the fibrotic activity of lung fibroblasts.

### ABHD17B and ABHD17C promote collagen expression in human HSCs

ABHD17B belongs to the ABHD17 family, which includes ABHD17A and ABHD17C. All three family members are expressed in primary human HSCs, as indicated by RNA sequencing data^12,13,15^. *Abhd17a* and *Abhd17b* are also induced in mouse HSCs with fibrotic injury *in vivo* (**Supplementary Fig. 8a**)^16^. These data prompted us to ask whether other ABHD17 family members also affect collagen expression in human HSCs. We found that depletion of *ABHD17B* and *ABHD17C* each reduced *COL1A1* expression in the three primary HSC donor lines tested, while depletion of *ABHD17A* was associated with modest reduction of *COL1A1* in only one line (**Fig. 2h**). In addition, depletion of *ABHD17B* did not affect *ABHD17A* or *ABHD17C* expression except for a small decrease in *ABHD17A* observed in one line. (**Supplementary Fig. 8b-d**). These results suggest that the phenotypes observed with depletion of *ABHD17B* are not affected by changes in expression of other ABHD17 family members.

### Analysis of ABHD17B structure

ABHD17 family proteins were identified as depalmitoylases in a screen for serine hydrolases that increase the turnover of palmitate^20^. ABHD17A has the strongest depalmitoylase activity, which was shown to be mitigated with a Ser to Ala mutation^20^. We performed structural analysis of ABHD17B with molecular dynamics (MD) simulations following *de novo* folding with AlphaFold^22^ or I-Tasser^60^ (**Supplementary Fig. 9a)**. The backbone Cα atoms stabilize with lower root mean square deviations (RMSD) for the structure generated with AlphaFold, therefore we used this structure for further analysis to define the active site of ABHD17B. Structural analysis of ABHD17B identified Ser170 on a loop between an alpha helix and beta sheet proximal to Asp235. His264 is located on a second loop between a different alpha helix and beta sheet, capable of forming a catalytic triad representative of alpha beta hydrolases (**Supplementary Fig. 9b, c**)^23^. These findings suggested that Ser170 has a high likelihood of being the active residue for depalmitoylation and that mutation of this residue would disrupt depalmitoylase activity. We performed a serine activity based probe assay (SABP) to quantify the activity of wildtype (WT) ABHD17B and ABHD17B containing a Ser to Ala mutation at residue 170 (S170A) and found that the mutation resulted in reduced hydrolase activity (**Fig. 3a**). These findings indicate that S170 is required for serine hydrolase/depalmitoylase activity in ABHD17B.

**Fig. 3.**
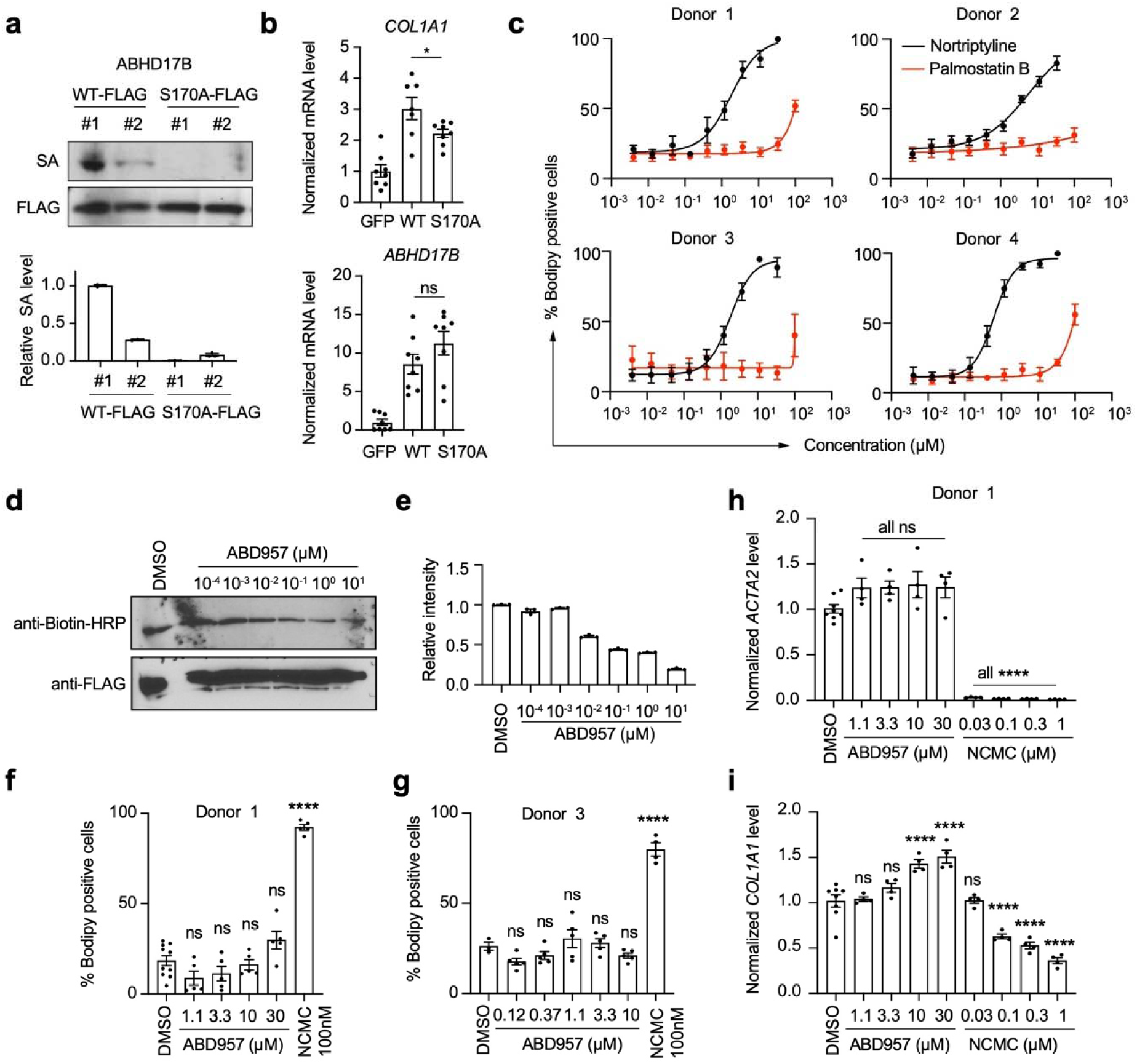
Depalmitoylation activity of ABHD17B is not required for HSC activation or collagen expression. (a) HEK-293T cells expressing wildtype (WT) ABHD17B-FLAG and ABHD17B-S170A-FLAG were lysed and treated with a desthiobiotin-labeled serine based activity probe (SABP). Serine activity was quantified by streptavidin (SA) HRP. Total protein was quantified by anti-FLAG antibody. Top: representative images. Bottom: each band was quantified three times using Fiji. Error bars represent mean ± SEM. Results are representative of two independent experiments. (b) *COL1A1* and *ABHD17B* were quantified by qRT-PCR in HSCs transduced with lentivirus expressing GFP, ABHD17B-WT or ABHD17B-S170A. Expression was normalized to *PSMB2*. Error bars represent mean ± SEM (n=8). ns indicates not significant and * indicates p<0.05 (2-tailed unpaired t-test between WT and S170A). Results are representative of two independent experiments. (c) HSCs were treated with nortriptyline or palmostatin B at indicated concentrations for 48 hr. Cells were fixed and stained with Bodipy and Hoechst. Each dot represents the averaged value of six replicates. Error bars represent mean ± SD (n=6). (d-e) HEK-293T cells expressing wildtype (WT) ABHD17B-FLAG and ABHD17B-S170A-FLAG were treated with increasing concentrations of ABD957. Cells were then lysed and treated with a desthiobiotin-labeled serine based activity probe (SABP) on M2 anti-FLAG beads. Serine activity was quantified by streptavidin HRP, and protein levels bound to M2 beads were quantified by probing with anti-FLAG antibody (d). Band intensities were quantified three times using Fiji and normalized to FLAG signal (e). Error bars represent mean ± SEM. Results are representative of two independent experiments. (f-g) HSCs from donor 1 (f) and 3 (g) were treated with ABD957 for 48 hr before Bodipy and Hoechst staining. Nanchangmycin (NCMC) served as a positive control^13^. Each dot represents one replicate. Error bars represent mean ± SEM (n≥5). ns indicates not significant and **** indicates p<0.0001 (one-way ANOVA test). (h-i) HSCs from donor 1 were treated with ABD957 or NCMC for 48 hr. *ACTA2* (h) and *COL1A1* (i) levels were quantified. Each dot represents one replicate. Error bars represent mean ± SEM (n≥4). ns indicates not significant and **** indicates p<0.0001 (one-way ANOVA test).

Next we asked how the S170A mutation affected gene expression in HSCs. Ectopic expression of ABHD17B-WT was associated with an increase in *COL1A1* expression, and this induction was blunted with expression of comparable levels of ABHD17B-S170A (**Fig. 3b**). This trend was also observed for expression of the fibrotic markers *TGFBR1* and *TIMP1,* while a significant change in *ACTA2* expression was not observed (**Supplementary Fig. 10a**). These results suggest that S170 is required for ABHD17B to fully promote fibrotic gene expression, but there may also be mechanisms that function independent of depalmitoylation.

### Depalmitoylase activity of ABHD17B is not required to promote HSC activation

To explore the role of depalmitoylation further, we treated HSCs with two depalmitoylase inhibitors. We treated HSCs with increasing concentrations of the inhibitor Palmostatin B (PalmB), which inhibits ∼80% of ABHD17B depalmitoylase activity at 10 µM^20^. A modest increase in lipid droplet formation was observed at the highest concentration (100 µM), while there was no difference at lower concentrations (**Fig. 3c**). We next evaluated the effect of the depalmitoylase inhibitor ABD957, which selectively inhibits the activities of ABHD17A, ABHD17B, ABHD17C, CES2 and ABHD6^24^. We found that the inhibitor also reduced the activity of ABDH17B by ∼80% at 10 µM (**Fig. 3d, e**); however, even at concentrations up to 30 µM, ABD957 had no effect on lipid accumulation (**Fig. 3f, g**) and did not reduce *ACTA2* or *COL1A1* expression (**Fig. 3h, i**). Similar results were observed when treating HSCs overexpressing ABHD17B with ABD957 (**Supplementary Fig. 10b**). These results show that neither global inhibition of depalmitoylase activity by PalmB nor more specific inhibition of the depalmitoylase activity of the ABHD17 family proteins by ABD957 affects the fibrotic activity of HSCs.

### Analysis of global gene expression and interacting partners

We next performed RNA-seq and differential expression analysis to define genes and pathways affected by depletion of *ABHD17B* that may not involve depalmitoylation. *ABHD17B* was depleted with two different siRNA duplexes, and each was compared to the nontargeting control (**Supplementary Table 7**). Results from both siRNAs were combined **(Supplementary Methods**), and depletion of *ABHD17B* was associated with a decrease in expression of genes involving the cytoskeleton, adhesion, contractility, ECM, and spindle formation (**Fig. 4a and Supplementary Fig. 11**). Heatmaps are shown for genes associated with these Gene Ontology (GO) categories to highlight mRNAs most affected by depletion of *ABHD17B* (**Fig. 4b and Supplementary Fig. 12**). These results suggest that ABDH17B promotes expression of genes regulating ECM production along with contractility and adhesion to drive fibrosis.

**Fig. 4.**
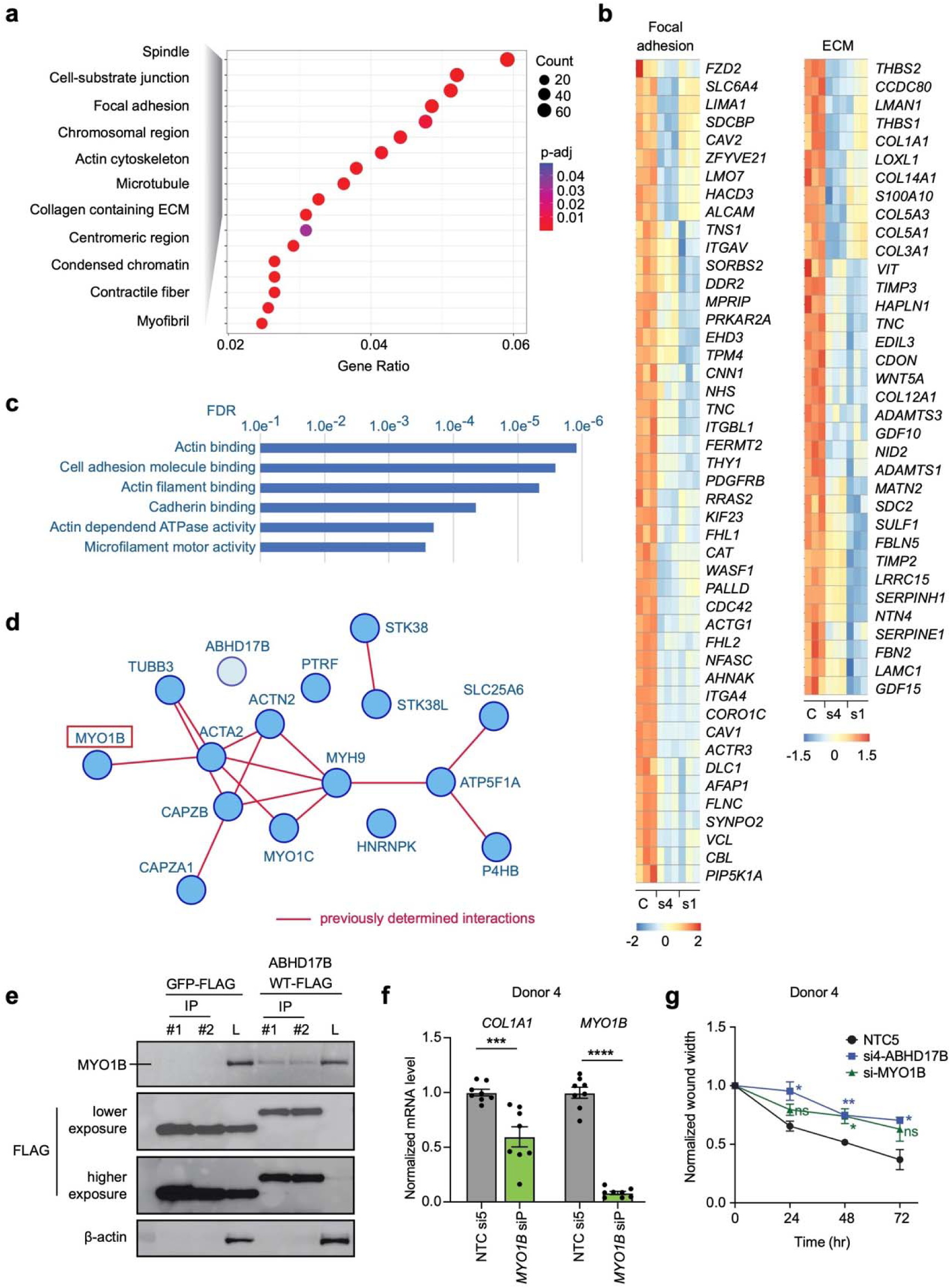
ABDH17B interacts with pathways involving contractility, adhesion, ECM, and the cytoskeleton. (a) RNA-seq and differential expression analysis were performed using nontargeting control (NTC) si5, and two siRNAs targeting *ABHD17B*. Dot plot displays the Gene Ontology (GO) terms most enriched in the genes repressed with depletion of *ABHD17B*. The color of each dot represents the adjusted p-value, and the size of the dot represents gene count. (b) Heatmaps show expression of the repressed gene set for indicated GO categories for HSCs transfected with NTC (C), siRNA4 targeting *ABHD17B* (s4) and siRNA1 targeting *ABHD17B* (s1). Expression is centered and scaled by row (gene). (c) Proteins that interact with ABHD17B were identified by precipitation of ABHD17B-FLAG and GFP-FLAG followed by mass spectrometry (MS) (**Supplementary Table 7**). Enrichment of GO categories (Molecular Functions) for the 15 proteins showing the strongest interaction with ABHD17B (String-db.org) is shown. (d) The interactions between the 15 proteins showing the strongest interaction with ABHD17B are displayed in addition to ABHD17B. Dark red lines indicate experimentally-determined interactions (String-db.org). (e) HSCs were transfected with lentivirus to express GFP-FLAG or ABDH17B-FLAG for 48 hr before anti-FLAG precipitation followed by probing for MYO1B (top) and FLAG expression (bottom). Two precipitations were performed for each condition (#1 and #2) and compared to total lysates (+L). Data is representative of two separate experiments. (f) Relative mRNA expression was quantified by qRT-PCR in primary human HSCs (Donor 4) treated witih non targeting siRNAs (NTC si5) and pooled siRNAs targeting *MYO1B*. Error bars represent mean ± SEM (n=8). *** indicates p < 0.001 and **** indicates p< 0.0001 (2-tailed unpaired t-test). Data are representative of three independent experiments. (g) Wound healing assay was performed in HSCs (Donor 4) transfected with indicated siRNAs. Normalized wound width was calculated at the indicated time points from three individual scratches. Error bars represent mean ± SEM (n=3). ns indicates not significant, (p=0.24 for si-MYO1B 24 hr, p=0.097 for si-MYO1B 72 hr), * indicates p < 0.05 and ** indicates p< 0.01 (one-way ANOVA test). Data are representative of two independent experiments.

We then evaluated the protein partners of ABHD17B. We ectopically expressed ABHD17B-FLAG and control GFP-FLAG in HSCs before performing precipitation and mass spectrometry (MS). Proteins showing the strongest enrichment with ABHD17B precipitation **(Supplementary Table 8, bold)** are associated with pathways (**Fig. 4c**) similar to those affected by depletion of *ABHD17B* (**Fig. 4a**). We next evaluated interactions among the 15 proteins showing the strongest interaction with ABHD17B (**Fig. 4d**). One large cluster is identified in which proteins are linked by experimental data supporting their interaction (red lines), while a second cluster contains the serine/threonine kinases STK38 and STK38L. ABH17B is also included in this analysis, but there was no previous evidence of interaction. Together, RNA-seq and MS data indicate that ABHD17B promotes expression of gene products involved in ECM production, contractility, and adhesion, and ABHD17B also directly interacts with proteins in these pathways. ABHD17 proteins localize to early endosome and the plasma membrane^20,30^ in a pattern similar to that observed for MYO1B^31^. Given this co-localization, we focused on the interaction predicted between ABHD17B and MYO1B. HSCs were transduced with lentivirus expressing *ABHD17B-FLAG* or *GFP-FLAG* followed by anti-FLAG immunoprecipitation and probing for MYO1B, which demonstrated enrichment of MYO1B protein with precipitation of ABDH17B (**Fig. 4e**).

We then depleted *MYO1B* in human HSCs to determine if the effect was similar to that observed with depletion of *ABHD17B*. Reduction in *MYO1B* was associated with attenuated expression of *COL1A1* (**Fig. 4f, Supplementaray Fig. 13a**), and depletion of either *ABDH17B* or *MYO1B* resulted in reduced migration in a wound healing assay (**Fig. 4g, Supplementary Fig. 13b**). Together, these results show that ABDH17B and MYO1B interact, and each promotes *COL1A1* expression and HSC migration, suggesting that they may work together to regulate these processes.

### Loss of Abhd17b protects against the development of liver fibrosis in vivo

*Abhd17b+/-* mice were intercrossed, and *Abhd17+/+, Abhd17+/-*, and *Abhd17b-/-* offspring were generated in expected Mendelian ratios (not shown). *Abhd17b-/-* mice bred normally, and liver histology appeared no different than that of *Abhd17b+/+* mice (**Fig. 5a**). Mice were treated with carbon tetrachloride (CCl_4_) to induce liver fibrosis, and *Abhd17b-/-* mice showed reduced fibrosis as measured by hydroxyproline (**Fig. 5b**) and collagen proportionate area (**Fig. 5c, d**). *Abhd17b-/-* mice also showed reduced expression of *Col1a1*, *Acta2*, *Tgfb1*, *Timp1*, and *Il1b* (**Fig. 5e-i**). Reduction in αSMA (encoded by *Acta2*) was also observed in *Abhd17b-/-* mice along with ductular reaction, as measured by Krt19 immunohistochemistry (IHC) (**Fig. 5j**). *Abhd17b-/-* mice also showed mild weight gain over four weeks of treatment while *Abhd17b+/+* did not gain weight (**Supplemenetary Fig. 14)**. These results suggest that *Abhd17b* is necessary for a robust fibrotic response in the setting of chronic liver injury.

**Fig. 5.**
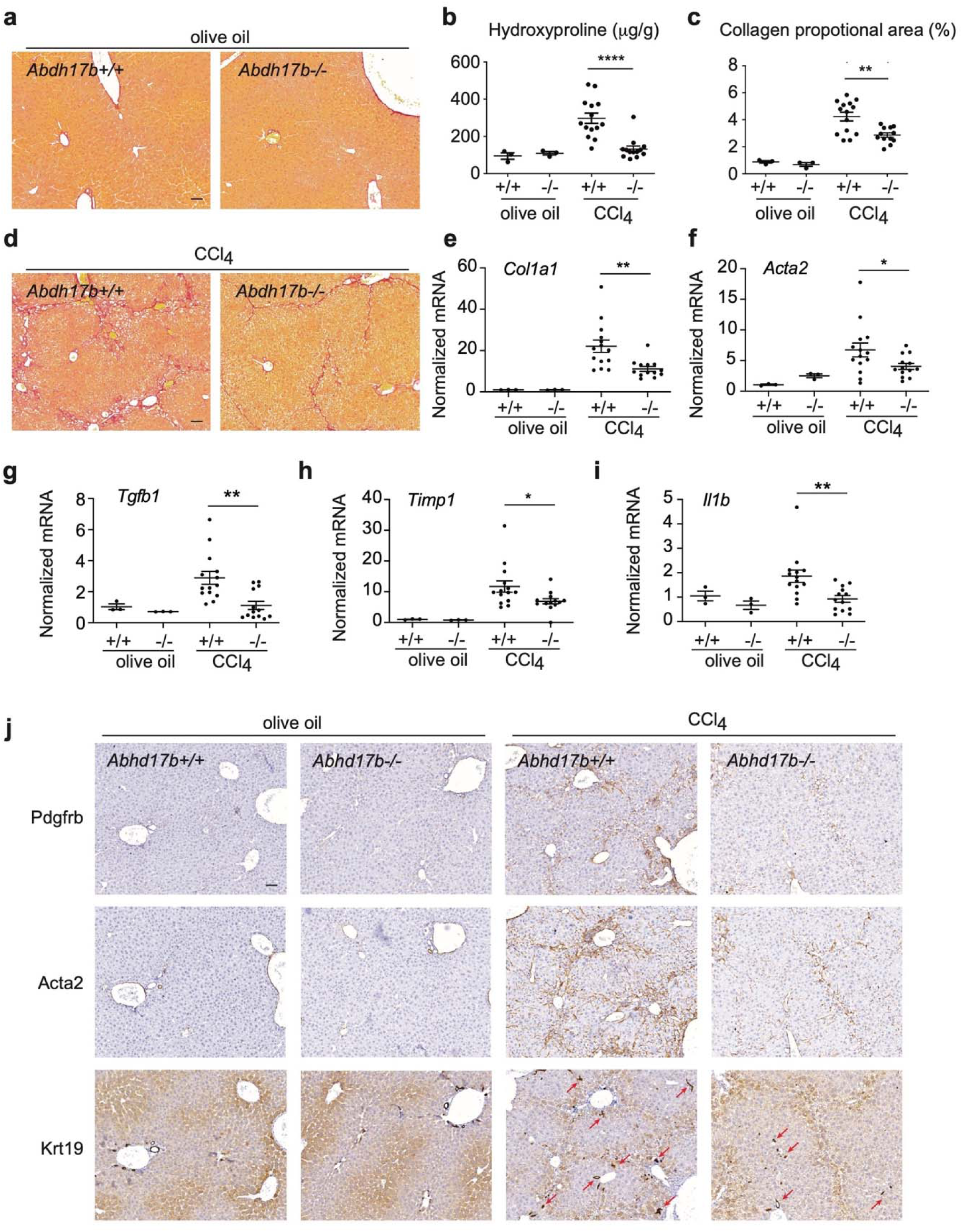
Loss of *Abhd17b* protects against development of liver fibrosis. (a) Representative Sirius red staining results of livers from *Abhd17b+/+* and *Abhd17b-/-* mice receiving control olive oil via oral gavage three times a week for four weeks. Scale bar indicates 50 μm. (b) Hydroxyproline analysis was performed using mouse liver samples. Error bars represent mean ± SEM (n=14, 13 for CCl_4_ treatment), **** indicates p<0.0001 (2-tailed unpaired t-test). (c) Collagen proportionate area (CPA) was calculated based on Sirius red staining. Error bars represent mean ± SEM (n=14, 12), ** indicates p<0.01 (2-tailed unpaired t-test). (d) Representative Sirius red staining results of livers from *Abhd17b+/+* and *Abhd17b-/-* mice recieing CCl_4_ via oral gavage three times a week for four weeks. Scale bar indicates 50 μm. (e-i) Relative mRNA expresion was analyzed for the indicated gene products by qRT-PCR from liver samples. Error bars represent mean ± SEM (n=14, 13), * indicates p<0.05 and ** indicates p<0.01 (2-tailed unpaired t-test). (j) Representative IHC staining for Pdgfrb, Acta2 and Krt19 in livers from *Abhd17b+/+* and *Abhd17b-/-* mice recieving either control olive oil or CCl_4_ via oral gavage three times a week for four weeks. Scale bar indicates 50 μm. Red arrows indicate examples of Krt19 positive cells.

## Discussion

Extent of fibrosis remains the strongest predictor of mortality in the setting of chronic liver disease^32^. While treating the underlying causes of liver injury have been successful in halting progression and even promoting regression of fibrosis^9,33^, effective treatments are not available for many sources of disease^2,3,34^. HSC activation and transdifferentiation into HSC myofibroblasts as a result of chronic injury is the primary process leading to accumulation of collagen and other components of the ECM that form the fibrotic scar^4–6^. Here, we performed an siRNA screen to identify gene products necessary to maintain the fibrotic phenotype of HSC myofibroblasts, with the goal of defining new therapeutic targets to inhibit fibrosis. The screen identified ABHD17B, a member of the ABHD17 family of proteins, which have depalmitoylase activity^20,24^. ABHD17 proteins accelerate palmitate turnover on PSD95 and N-Ras^20^, and ABD957, an inhibitor of ABHD17 depalmitoylase activity, impairs N-Ras depalmitoylation in human acute myeloid leukemia cells^24^. ABHD17B was also reported to attenuate TGF-β-induced palmitoylation of hexokinase 1 (HK1) in LX-2 cells, leading to reduced HK1 secretion^35^. From these studies, it was not clear if and how ABHD17B is involved in HSC activation and liver fibrosis, prompting further investigation.

Through structural and mutation analysis we find that Ser 170 is the active residue for protease/depalmitoylase activity of ABDH17B. While we do not find depalmitoylase activity to be required for its profibrotic function, palmitoylation itself could still play a role in ABHD17B function, as its N-terminal region is itself palmitoylated to mediate membrane interactions^20,30^ and could affect binding to other proteins at membrane surfaces, including MYO1B^31^.

RNA-seq and MS studies were performed to better understand how ABHD17B may function independent of depalmitoylase activity. Depletion of ABHD17B affects expression of genes that control ECM production, adhesion, contractility, and the cytoskeleton, while MS studies revealed that ABHD17B associates with proteins involved in cell adhesion/cadherin binding and microfilament motor activity linked to actin in a network that includes MYO1B and MYO1C (**Fig. 4d**), MYO1B also regulates expression of *COL1A1* and HSC migration (**Fig. 4f, g**), and *Myo1c*-deficient mice are protected from liver fibrosis^36^, suggesting that controlling the activity of type I myosins is one mechanism by which ABDH17B may regulate the activity of HSCs.

Additional interacting partners identified by MS could also contribute to similar pathways. CAVIN1/PTRF is not part of any cluster, but in addition to a role in transcription terminiation^37^, it is also part of the caveolin complex^38^, which regulates actin cytoskeleton and adhesion in HSCs^39^. Multiple nuclear functions are attributed to HNRPNK^40,41^, but it also associates with calponin^42^, a protein that binds actin in a complex regulating contractility^43^. STK38 and STK38L (NDR1/2) are serine-threonine kinases that form a separate cluster. These proteins are related to LATS1/2 in the HIPPO pathway^44^, and STK38 regulates MHY6 and sarcomere assembly in cardiomyocytes^45^, suggesting that it could modulate other non-muscle myosins in fibroblasts. The significance of interaction with mitochondrial proteins (ATP5F1A) is unclear but it is linked to MHY9 (**Fig. 4d**).

Environmental stiffness regulates the actin cytoskeleton and collagen expression in fibroblasts^46^. The interacting partners identified by MS, and confirmed with additional analysis, in the case of MYO1B, suggest ABHD17B may promote the fibrotic activity of HSCs by mediating the dynamic response to signals of environmental stiffness transmitted to the cystoskeleton through adhesion molecules and interpreted by type I myosin molecules. Type I myosins can sense tension on actin filaments^47,48^, and interactions with ABHD17B on membrane surfaces may modulate myosin-mediated endosomal transport^49,50^, including glucose transporters and TGF-β receptors that promote fibrosis^17,51–54^.

In summary, we performed an siRNA screen to identify genes that could be targeted to promote inactivation of HSCs, leading to the identification of ABHD17B. Depletion of *ABHD17B* in primary HSCs decreased collagen expression and modulated pathways involving the actin cytoskeleton, contractility, focal adhesion, and ECM. Furthermore, mice deficient in *Abhd17b* are protected from liver fibrosis. While we could not exclude the possible impact from *Abhd17b* depletion in other cell types from these *in vivo* experiments, the antifibrotic phenotype of *abhd17b*-deficient mice should be mediated at least partially via HSC inactivation, as demonstrated by our *in vitro* experiments with primary HSCs. Together, these findings suggest that targeting ABHD17B-dependent pathways in HSCs may be a viable approach to reduce fibrosis. Current small molecules inhibiting the depalmitoylase activity of ABHD17B do not affect this process, and further studies will be required to develop approaches to inhibit the activity of ABHD17B.

## Materials and methods

### Animals

All mouse experiments were approved by the IACUC of the Massachusetts General Hospital (2017000074). *Abhd17b+/-* (*C57BL/6N-Abhd17b^em^*^1^*^(IMPC)J^/Mmucd*) mice were purchased from the Mutant Mouse Resource and Research Center (MMRRC) and were maintained on C57BL/6 background. Male mice received 40% carbon tetrachloride (CCl_4_) diluted in olive oil or olive oil control by oral gavage (100 μl total volume) three times a week for four weeks^55^. *Abhd17b-/-* and *Abhd17b+/+* mice were generated from *Abhd17b-/-* and *Abhd17b+/+* siblings produced from *Abhd17b+/-* parents. Studies were initiated on age matched *Abhd17b-/-* and *Abhd17b+/+* mice at 8-9 weeks of age.

### Cell culture and compound treatments

Primary human hepatic stellate cells (HSCs) were purchased from Lonza and Zen-Bio and cultured as described^13^. Donor information is provided below. There is limited availability of samples from individual donors. The cells used for the screening (Donors 1-4, Lonza) were from donated tissues of deceased donors with consent or legal authorization. The screen was conducted using primary HSCs at passage 8. Cells at passages 8-10 were used for the other expriments.

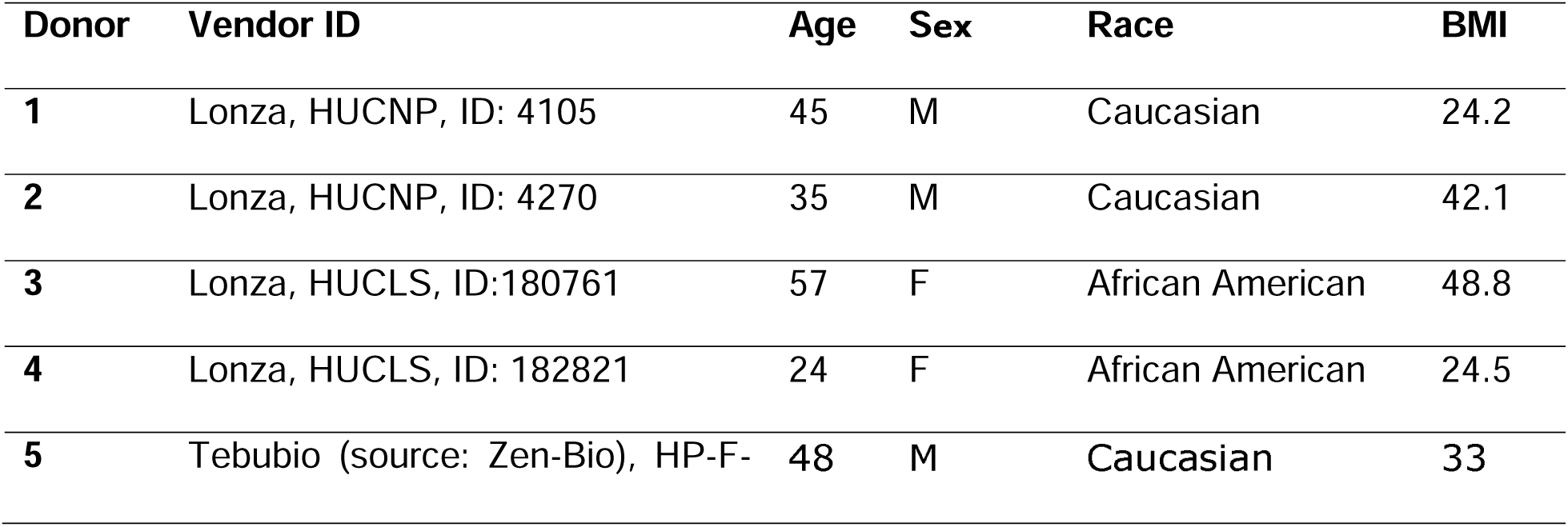
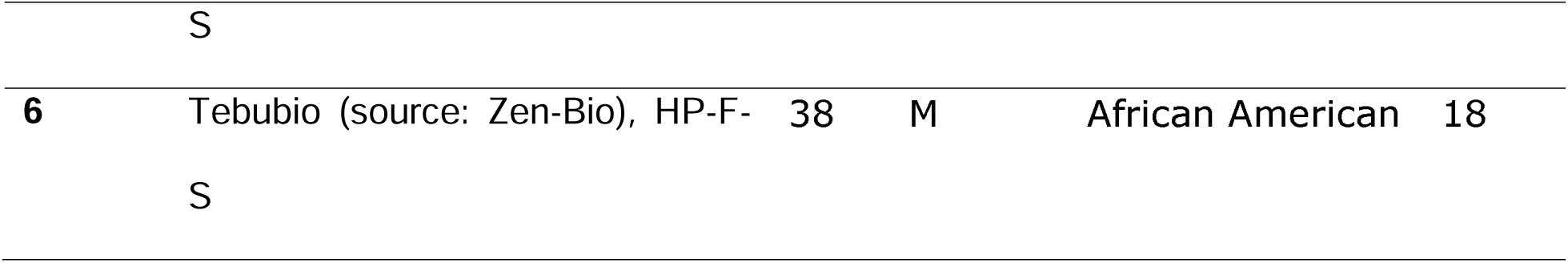

Primary lung fibroblasts were collected from deidentified discarded excess tissue from a healthy donor (699) through the MGH Fibrosis Translational Research program and cultured as described previously^56^. HEK-293 cells were obtained from ATCC (CRL-1573) and cultured in DMEM supplemented with 10% fetal calf serum and 1% Penicillin/Streptomycin.

Palmostatin B was purchased from Calbiochem (Cat. # 178501). ABD957 was synthesized by chemists from Boehringer Ingelheim (purity: 93% as determined by HPLC-MS; MS (ESI^+^): m/z = 628.6 [M+H]^+^, 628.2 calculated). Nanchangmycin was purchased from Adooq (Cat. # A10621). Nortriptyline was purchased from Sigma (Cat. # N7261). HSCs were treated with compounds for 48hr at concentrations as indicated before fixed or lysed.

### Reverse transfection of siRNAs in the screen and follow-up experiments

For the screening, the transfection was performed in 384-well plates (Corning, 3764) in a high-throughput manner using Bravo Automated Pipettor (Agilent) and Multidrop Combi Reagent Dispenser (Thermo Fisher Scientific). For each well in a 384-well plate, 1.25 µL siRNAs (in 1 µM stocks) were diluted in 8.5 µL Opti-MEM (to make the final concentration of 25 nM) in the wells, and then 10 µL diluted Dharmafect-1 (Horizon Discovery, T-2001-02, 9.95 µL Opti-MEM + 0.05 µL Dharmafect-1) was added. After a 40min incubation, 750 HSCs in 30 µL antibiotic-free transfection media (DMEM + 16% FBS) were added to the siRNA and Dharmafect-1 mixture in each well. Cells were fixed and stained with Bodipy and Hoechst 72 hr after transfection. Cells-to-CT 1-Step Taqman Kit was used to quantify mRNA expression. *ACTA2*, *COL1A1* and target mRNA were each quantified in the same well with the endogenous control *PSMB2.* The gene-specific TaqMan Real-time PCR Assays for qPCR analysis are provided in supplmentary materials and methods.

The siRNAs used in the screen and the siRNAs targeting *FAM160B2*, *MGAT5B*, *TMEM134*, *TTTY2* and *TTY2B* were provided by the Institute of Chemistry and Cell Biology (ICCB)- Longwood screening facility, and the relevant information is provided in the supplementary materials and methods and supplementary tables.

For the validation experiments, the transfection in HSCs was scaled up to 96-well plates (for staining) and 12-well plates (for qRT-PCR) proportionally based on surface area from the protocol described above. For transfection of siRNAs in primary lung fibroblasts, cells were reverse transfected using lipofectamine 2000 (Thermo Fisher Scientific, 11668019) according to manufacturer’s manual at 5000 cells/cm^2^ with 20 nM siRNA. Cells were lysed 72 hr after transfection. The siRNAs used in the validation experiments were purchased from Horizon Discovery as listed in supplementary materials and methods.

### Analysis of α-SMA fibers and extracellular collagen deposition (scar-in-a-jar assay)

Primary human HSCs were seeded in 384-well plates (CellCarrier-384 Ultra plates, Cat. # 6057300) in stellate cell medium (ScienCell/Innoprot, Cat. # 5301-b) supplemented with 2% FCS (Thermo Fisher Scientific), 1% penicillin/streptomycin, and stellate cell growth supplement (ScienCell/Innoprot, Cat. # 5352), and transfected using JetPrime transfection reagent (Polyplus, Cat. # 101000027). The final siRNA concentration per well was 16 nM with 0.8 µl transfection reagent per well. Cells were serum starved the next day, followed by stimulation with TGF-β (10 ng/ml, R&D, Cat. # 240-B) in conditions of molecular crowding (37.5 mg/ml Ficoll 70 and 25 mg/ml Ficoll 400, GE Healthcare, Cat. # 17-0310-10 and # 17-0300-10) and vitamin C supplementation (0.2 mM; Sigma, Cat. # A8960) to promote collagen deposition. 72 hr post TGF-β stimulation, cells were fixed in 80% Methanol for 30 min, washed in PBS, and permeabilized using 1% TritonX 100 in PBS. Cells were blocked in 3% BSA for 30 min at RT and incubated with anti-alpha-SMA antibody (1:1000; Sigma, Cat. # A2547) and anti-collagen type I antibody (1:1000; Sigma, Cat. # SAB4200678) in PBS at 37° C for 1.5 hr. Cells were washed with PBS and incubated with secondary antibodies for 30 min at 37° C (1:1000 AF568 goat anti-mouse IgG1, Thermo Fisher Scientific, Cat. # A-21124; and 1:1000 AF647 goat anti-mouse IgG2a, Thermo Fisher Scientific, Cat. # A-21241). Nuclei were stained with Hoechst (1 µM, Molecular Probes, Cat. # H-3570), cells with HSC Cellmask GreenTM at 0.2 μg/ml (Invitrogen, Cat. # H32714). Nuclear, cytoplasm, αSMA and collagen-I images were acquired using the IN Cell Analyzer 2200, images transferred to Perkin Elmer’s Columbus Image Storage and Analysis system, and analyzed using a custom image analysis protocol with total cell number, number of α-SMA fibers/cell, and collagen area/total cell number being quantified, as described^57^.

### Lentiviral transduction of HSCs

Lentivirus was produced in HEK-293T cells by transfecting plasmids expressing either EGFP_FLAG, ABHD17B-FLAG, or S170A-ABHD17B-FLAG (nucleotide sequences for ABHD17B and S170A are provided in **Supplementary Fig. 15**) along with pMD2.G (Addgene plasmid # 12259) and psPAX2 (Addgene plasmid # 12260) using X-tremeGENE 9 transfection reagent^13^. Lentivirus was administered with 10 μg/mL polybrene (Sigma-Aldrich, Cat. # TR-1003-G) for 24 hr before the first media change. After 24 hr, cells were selected with puromycin (1 µg/mL, Cat. # A1113803) for 4 days. RNA was harvested in Trizol, and cell lysates for Western blot were harvested in 1μg/ml Pierce IP Lysis buffer.

### Serine activity probe based assays

Thermo Scientific™ ActivX™ Desthiobiotin-FP Serine Hydrolase Probe was purchased from Thermo Fisher (Cat. # 88317). Anti-FLAG® M2 Magnetic Beads (Cat. # M8823-1ML) were purchased from Milipore Sigma. Blots were developed with either anti-biotin, HRP-linked antibody (Cell Signaling, Cat. # 7075), streptavidin (SA) HRP (Cell Signaling, Cat. # 3999) or anti-FLAG antibody (Cell Signaling, Cat. # 14793). HEK-293T cells were transiently transfected using X-tremeGENE™ 9 DNA Transfection Reagent (Roche, Cat. # XTG9-R0) with 6 µL of transfection reagent to 2 µg plasmid for 48 hr. Cells were harvested in 300 µL Pierce™ IP Lysis Buffer (Thermo Scientific, Cat. # 87787) and frozen at −80° C. Lysates were cleared at 10,000 *g,* and an equal amount of equilibrated Millipore Sigma Anti-FLAG^®^ M-2 Magnetic Beads (Cat. # M8823) in Pierce Lysis Buffer was added, followed by 2 washes in 1 mL Pierce Lysis Buffer using magnetic rack. Beads were resuspended in 50 μL of Pierce Lysis Buffer and labeled with addition of 1 µL of ActivX™ Desthiobiotin-FP stock (at 100 µM in DMSO) for 30 min on bead, rocking at RT with occasional vortexing. NuPAGE™ Sample Reducing Agent (10X) (Invitrogen, Cat. # NP0009) and NuPAGE™ LDS Sample Buffer (4X) (Invitrogen, Cat. # NP 0007) were added to samples before boiling at 95 °C for 5 min. Samples were loaded on Bolt™ 4 to 12%, Bis-Tris, 1.0 mm, Mini Protein Gels (Cat. # NW04120BOX) in NuPAGE™ MOPS SDS Running Buffer (20X) (Cat. # NP 0001002). SeeBlue™ Plus2 Pre-stained Protein Standard (Cat. # LC5925) was used as a molecular weight marker. After electrophoresis, material was transferred to iBlot™ 2 Transfer Stacks, nitrocellulose (Cat. # IB23001) using an iBlot™ 2 Gel Transfer Device (Cat. # IB21002S). Nitrocellulose was blocked in 1% BSA (Thermo Scientific, Cat. # 37520) and probed using anti-biotin antibody or SA-HRP overnight at 4 °C. Blots were washed 3 times with Tris Buffered Saline-Tween (TBST) buffer (Boston BioProducts, Cat. # IBB-181–6) and developed using SuperSignal™ West Pico PLUS Chemiluminescent Substrate (Thermo Scientific, Cat. # 34580). Blots were then stripped according to manufacturer’s specifications with Thermo Fisher Scientific Restore™ Western Blot Stripping Buffer (Cat. # 21059), washed in TBST 3 times, re-blocked in 1% BSA and re-probed with anti-FLAG antibody overnight at 4° C. Nitrocellulose was then washed 3 times in TBS-T and probed with goat-anti-anti-Rabbit IgG (H+L) secondary antibody, HRP (Invitrogen, Cat. # 34260).

### qRT-PCR analysis

RNA samples were extracted using TRIzol (Invitrogen, cat# 15596026). Using 1 μg total RNA as input, reverse transcription was performed with the iScript gDNA Clear cDNA Synthesis Kit (BIO-RAD, 1725035) according to manufacturer’s instructions. TaqMan Universal PCR Master Mix (Applied Biosystems, cat# 4305719) and TaqMan Real-time PCR Assays (ThermoFisher Scientific) were used for the quantitative real-time PCR analysis of the cDNA samples. For each qRT-PCR experiment, each dot represents the mean value for a single sample analyzed in duplicate to quadruplicate and PSMB2 as the endogenous control^13^. The gene-specific TaqMan Real-time PCR Assays used in this study are listed in supplementary materials and methods.

The sequences of the primers used for analyzing the overexpression level of wildtype and mutant *ABHD17B* are 5’-ggactgaagatgaagtcattgacttttcacatgg-3’ (forward), and 5’-cttatcgtcgtcatccttgtaat cg-3’ (reverse) and were selected to recognize both *ABHD17B-WT* and *ABHD17B-S170A* transcripts.

For quantification of siRNA-mediated silencing in the scar-in-a-jar assay, cells were transfected and treated as described, lysed in RLT Plus buffer, and RNA isolated with the RNeasy Plus Kit according to the manufacturer’s protocol (Qiagen, Cat. # 74134). To obtain sufficient material for qRT-PCR, lysates from 6-12 wells were pooled prior to RNA isolation. 150 ng RNA were reverse transcribed using the High-capacity cDNA Archive Kit (Applied Biosystems, Cat. # 4322169) and Taqman assays performed using TaqMan Universal PCR Master Mix (Applied Biosystems, Cat. # 4318157), probes listed in the table above, and the ViiA 7 Real-Time PCR System (Thermo Fisher Scientific).

### Precipitation and mass spectrometry

Precipitation experiments were performed following transduction of HSCs with lentivirus expressing ABHD17B-FLAG or GFP-FLAG. Cells were selected with puromycin (1 μg/ml) for 4 days and grown to confluence before harvest in Pierce Lysis Buffer. Lysates were frozen at −80° C, thawed at 4° C, and cleared by centrifugation at 4500 *g* for 10 min at 4° C. Lysates were incubated with anti-FLAG (M2) beads on a rocker for 30 min at 4° C with vortexing at 10 min intervals. Beads were washed two times with 1 mL Pierce Lysis Buffer using a magnetic rack and eluted in 100 μl Pierce lysis buffer containing 3X FLAG® Peptide (Sigma, Cat. # F4799) at 20 μM. Eluted proteins were boiled at 95° C, and gel electrophoresis was performed. For MS experiments, after electrophoresis, proteins were visualized with Coomassie Blue. Bands were excised above and below the location of GFP protein for MS analysis at the Taplin Mass Spectrometry following approaches previously described^58^. Data from two biological experiments, each in duplicate were analyzed. DESeq2^59^ was applied to spectral counts to obtain a list of proteins showing differential abundance between ABDH17B IP samples and control GFP IP samples. For Western pull down experiments gels were immediately transferred to iBlot™ 2 Transfer Stacks, and probed as in Serine Activity Assay using HRP chemiluminescence secondary antibodies where anti-FLAG antibody (Cell Signaling, Cat. # 14793) or anti-Myo1b antibody (abcam, Cat. # 194356) were used as the primary antibody.

### Protein structure prediction, molecular dynamics based analysis of ABHD17B

Full-length human ABHD17B structures were obtained either from deposit of *de novo* Deepmind Alphafold Structure Database^25^ or prepared *de novo* through I-Tasser^60^. All models of full-length ABHD17B structure were prepared in Schrödinger Maestro and minimized using Schrodinger Protein Preparation Wizard using OPLS2005^26^. Minimized structures were exported in .pdb format and prepared using solution builder from CHARMM-GUI^61^ with a salt concentration of 150 mM NaCl^62^. Molecular dynamics simulations were performed using NAMD 2.12^29^ with the CHARMM36m force field^63^ on full-length ABHD17B generated from either i-Tasser^64^ or AlphaFold2^22^. Visualization and analyses were performed using VMD 1.9.3^28^. The system was equilibrated for 10 ns restraining the Cα atoms of the protein (1.0 kcal/mol/A2) to allow for solvation. Root mean square deviations (RMSD) using backbone Cα atoms were calculated using VMD RMSD trajectory tool and exported and plotted using Graphpad Prism version 9. Three-dimensional representations were generated in Schrödinger ^26^.

### Wound healing analysis

HSCs were transfected with nontargeting control siRNAs and those targeting *ABHD17B* or *MYO1B* as described above using confluent cells split 1:1: One day after transfection, a p-20 pipette tip was used to generate a wound field. Cells were monitored and the width of the wounds were visualized via microscopy at indicated times. Images were taken using EVOS FL microscope and exported into ImageJ to measure relative wound width.

## Acknowledgements

We thank the staff at the ICCB-Longwood Screening Facility, and especially Richard Siu for their assistance. We thank Nicola Zimmermann and Sabine Weigle at Boehringer Ingelheim for conducting *in vitro* analysis of α-SMA fibers and extracellular collagen deposition. We thank members of the Mullen lab and colleagues at Boehringer Ingelheim, including John Broadwater, Peter Seither, and Christofer Tautermann for helpful discussion, as well as the Taplin Mass Spectrometry Core at Harvard Medical School. This work was funded through a grant from Boehringer Ingelheim Pharma GmbH Co KG. The graphic abstract contains images from Servier Medical Art (https://smart.servier.com/) and BioRender (https://biorender.com).

## Authors’ Contributions

W.L. and A.C.M. conceived the study with C.B-K., J.F.R., J.F.D, and D.M.S. W.L., R.P.S., and C.S. performed the experiments with assistance from J.Y.C., J.S., S.P.M., M.S., Y.Y., B.J.T., and M.L.B., and D.W. supported processing of imaging data. Protein structural analysis was performed by R.P.S. with support from A. W. and G. A., and G.A. also oversaw synthesis and analysis of ABD957. All other computational analyses were performed by L.P., R.K., and V. B. under the direction of S.J.H.S., and Z.L. and P.Z. with support from C.Z. The manuscript was written by W.L., R.P.S., C.S., D.M.S, and A.C.M. with input from all other authors.

## References

1. Poynard, T. et al. Impact of pegylated interferon alfa-2b and ribavirin on liver fibrosis in patients with chronic hepatitis C. Gastroenterology 122, 1303–1313 (2002).

2. Musso, G., Gambino, R., Cassader, M. & Pagano, G. A meta-analysis of randomized trials for the treatment of nonalcoholic fatty liver disease. Hepatology 52, 79–104 (2010).

3. Karlsen, T. H., Folseraas, T., Thorburn, D. & Vesterhus, M. Primary sclerosing cholangitis â€“ a comprehensive review. Journal of Hepatology 67, 1298–1323 (2017).

4. Friedman, S. L., Roll, F. J., Boyles, J. & Bissell, D. M. Hepatic lipocytes: the principal collagen-producing cells of normal rat liver. Proceedings of the National Academy of Sciences 82, 8681–8685 (1985).

5. Maher, J. J. & McGuire, R. F. Extracellular matrix gene expression increases preferentially in rat lipocytes and sinusoidal endothelial cells during hepatic fibrosis in vivo. Journal of Clinical Investigation 86, 1641–1648 (1990).

6. Mederacke, I. et al. Fate tracing reveals hepatic stellate cells as dominant contributors to liver fibrosis independent of its aetiology. Nature Communications 4, 1–11 (2013).

7. Friedman, S. L. Hepatic Stellate Cells: Protean, Multifunctional, and Enigmatic Cells of the Liver. 88, 125–172 (2008).

8. Bataller, R. & Brenner, D. A. Liver fibrosis. Journal of Clinical Investigation 115, 209–218 (2005).

9. Falize, L. et al. Reversibility of hepatic fibrosis in treated genetic hemochromatosis: A study of 36 cases. Hepatology 44, 472–477 (2006).

10. Kisseleva, T. et al. Myofibroblasts revert to an inactive phenotype during regression of liver fibrosis. Proceedings of the National Academy of Sciences 109, 9448–9453 (2012).

11. Troeger, J. S. et al. Deactivation of Hepatic Stellate Cells During Liver Fibrosis Resolution in Mice. Gastroenterology 143, 1073–1083.e22 (2012).

12. Chen, J. Y. et al. Tricyclic Antidepressants Promote Ceramide Accumulation to Regulate Collagen Production in Human Hepatic Stellate Cells. Scientific Reports 7, 44867–13 (2017).

13. Li, W. et al. Nanchangmycin regulates FYN, PTK2, and MAPK1/3 to control the fibrotic activity of human hepatic stellate cells. Elife 11, e74513 (2022).

14. Johnston, S. M., Shamu, C. E. & Smith, J. A. Automation Considerations for RNAi Library Formatting and High Throughput Transfection. eBook. eISBN 978-1-60805-940-9. Frontiers in RNAi 1, 21–39 (2014).

15. Zhou, C. et al. Long noncoding RNAs expressed in human hepatic stellate cells form networks with extracellular matrix proteins. Genome Medicine 1–20 (2016) doi:10.1186/s13073-016-0285-0.

16. Yang, W. et al. Single-Cell Transcriptomic Analysis Reveals a Hepatic Stellate Cell– Activation Roadmap and Myofibroblast Origin During Liver Fibrosis in Mice. Hepatology 74, 2774–2790 (2021).

17. Qi, Z., Atsuchi, N., Ooshima, A., Takeshita, A. & Ueno, H. Blockade of type β transforming growth factor signaling prevents liver fibrosis and dysfunction in the rat. Proc. Natl. Acad. Sci. 96, 2345–2349 (1999).

18. Dooley, S. et al. Smad7 prevents activation of hepatic stellate cells and liver fibrosis in rats. Gastroenterology 125, 178–191 (2003).

19. Tanahashi, N. et al. Chromosomal Localization and Immunological Analysis of a Family of Human 26S Proteasomal ATPases. Biochem. Biophys. Res. Commun. 243, 229–232 (1998).

20. Lin, D. T. S. & Conibear, E. ABHD17 proteins are novel protein depalmitoylases that regulate N-Ras palmitate turnover and subcellular localization. eLife 4, e11306 (2015).

21. Wynn, T. A. Integrating mechanisms of pulmonary fibrosis. J. Exp. Med. 208, 1339–1350 (2011).

22. Jumper, J. et al. Highly accurate protein structure prediction with AlphaFold. Nature 596, 583–589 (2021).

23. Rauwerdink, A. & Kazlauskas, R. J. How the Same Core Catalytic Machinery Catalyzes 17 Different Reactions: the Serine-Histidine-Aspartate Catalytic Triad of α/β-Hydrolase Fold Enzymes. Acs Catal 5, 6153–6176 (2015).

24. Remsberg, J. R. et al. ABHD17 regulation of plasma membrane palmitoylation and N-Ras-dependent cancer growth. Nat Chem Biol 17, 856–864 (2021).

25. Varadi, M. et al. AlphaFold Protein Structure Database: massively expanding the structural coverage of protein-sequence space with high-accuracy models. Nucleic Acids Res 50, D439– D444 (2021).

26. Schrödinger Release 2024-2: Maestro, Schrödinger, LLC, New York, NY, 2024.

27. Banks, J. L. et al. Integrated Modeling Program, Applied Chemical Theory (IMPACT). J. Comput. Chem. 26, 1752–1780 (2005).

28. Humphrey, W., Dalke, A. & Schulten, K. VMD: Visual molecular dynamics. J Mol Graphics 14, 33–38 (1996).

29. Phillips, J. C. et al. Scalable molecular dynamics with NAMD. J. Comput. Chem. 26, 1781– 802 (2005).

30. Martin, B. R. & Cravatt, B. F. Large-scale profiling of protein palmitoylation in mammalian cells. Nat. Methods 6, 135–138 (2009).

31. Almeida, C. G. et al. Myosin 1b promotes the formation of post-Golgi carriers by regulating actin assembly and membrane remodelling at the trans-Golgi network. Nat. Cell Biol. 13, 779– 789 (2011).

32. Hagström, H. et al. Fibrosis stage but not NASH predicts mortality and time to development of severe liver disease in biopsy-proven NAFLD. J. Hepatol. 67, 1265–1273 (2017).

33. Bonis, P. A. L., Friedman, S. L. & Kaplan, M. M. Is Liver Fibrosis Reversible? New Engl J Medicine 344, 452–454 (2001).

34. Sanyal, A. J. et al. Pioglitazone, vitamin E, or placebo for nonalcoholic steatohepatitis. New Engl J Medicine 362, 1675–85 (2010).

35. Chen, Q. et al. HK1 from hepatic stellate cell–derived extracellular vesicles promotes progression of hepatocellular carcinoma. Nat. Metab. 4, 1306–1321 (2022).

36. Arif, E. et al. Targeting myosin 1c inhibits murine hepatic fibrogenesis. Am. J. Physiol.-Gastrointest. Liver Physiol. 320, G1044–G1053 (2021).

37. Jansa, P., Mason, S. W., Hoffmann-Rohrer, U. & Grummt, I. Cloning and functional characterization of PTRF, a novel protein which induces dissociation of paused ternary transcription complexes. Embo J 17, 2855–2864 (1998).

38. Hill, M. M. et al. PTRF-Cavin, a Conserved Cytoplasmic Protein Required for Caveola Formation and Function. Cell 132, 113–124 (2008).

39. Ilha, M. et al. Exogenous expression of caveolin-1 is sufficient for hepatic stellate cell activation. J. Cell. Biochem. 120, 19031–19043 (2019).

40. Expert-Bezançon, A., Caer, J. P. L. & Marie, J. Heterogeneous Nuclear Ribonucleoprotein (hnRNP) K Is a Component of an Intronic Splicing Enhancer Complex That Activates the Splicing of the Alternative Exon 6A from Chicken β-Tropomyosin Pre-mRNA*. J Biol Chem 277, 16614–16623 (2002).

41. Takimoto, M. et al. Specific binding of heterogeneous ribonucleoprotein particle protein K to the human c-myc promoter, in vitro. J Biol Chem 268, 18249–18258 (1993).

42. Laury-Kleintop, L. D., Tresini, M. & Hammond, O. Compartmentalization of hnRNP-K during cell cycle progression and its interaction with calponin in the cytoplasm. J. Cell. Biochem. 95, 1042–1056 (2005).

43. Reynaert, H. Hepatic stellate cells: role in microcirculation and pathophysiology of portal hypertension. Gut 50, 571–581 (2002).

44. Zhang, L. et al. NDR Functions as a Physiological YAP1 Kinase in the Intestinal Epithelium. Curr Biol 25, 296–305 (2015).

45. Liu, J. et al. Stk38 Modulates Rbm24 Protein Stability to Regulate Sarcomere Assembly in Cardiomyocytes. Sci Rep-uk 7, 44870 (2017).

46. Mochitate, K., Pawelek, P. & Grinnell, F. Stress relaxation of contracted collagen gels: Disruption of actin filament bundles, release of cell surface fibronectin, and down-regulation of DNA and protein synthesis. Exp Cell Res 193, 198–207 (1991).

47. Laakso, J. M., Lewis, J. H., Shuman, H. & Ostap, E. M. Myosin I Can Act As a Molecular Force Sensor. Science 321, 133–136 (2008).

48. Greenberg, M. J., Lin, T., Goldman, Y. E., Shuman, H. & Ostap, E. M. Myosin IC generates power over a range of loads via a new tension-sensing mechanism. Proc. Natl. Acad. Sci. 109, E2433–E2440 (2012).

49. Raposo, G. et al. Association of Myosin I Alpha with Endosomes and Lysosomes in Mammalian Cells. Mol. Biol. Cell 10, 1477–1494 (1999).

50. Brandstaetter, H., Kendrick-Jones, J. & Buss, F. Myo1c regulates lipid raft recycling to control cell spreading, migration and Salmonella invasion. J. Cell Sci. 125, 1991–2003 (2012).

51. Bose, A. et al. Glucose transporter recycling in response to insulin is facilitated by myosin Myo1c. Nature 420, 821–824 (2002).

52. Chandrashekaran, V. et al. Purinergic receptor X7 mediates leptin induced GLUT4 function in stellate cells in nonalcoholic steatohepatitis. Biochim. Biophys. Acta (BBA) - Mol. Basis Dis. 1862, 32–45 (2016).

53. Chung, C. et al. Pentachloropseudilin Inhibits Transforming Growth Factor-β (TGF-β) Activity by Accelerating Cell-Surface Type II TGF-β Receptor Turnover in Target Cells. ChemBioChem 19, 851–864 (2018).

54. Chung, C.-L., Tai, S.-B., Hu, T.-H., Chen, J.-J. & Chen, C.-L. Roles of Myosin-Mediated Membrane Trafficking in TGF-β Signaling. Int. J. Mol. Sci. 20, 3913 (2019).

55. Alsamman, S. et al. Targeting acid ceramidase inhibits YAP/TAZ signaling to reduce fibrosis in mice. Science translational medicine 12, eaay8798 (2020).

56. Ramos, C. et al. Fibroblasts from Idiopathic Pulmonary Fibrosis and Normal Lungs Differ in Growth Rate, Apoptosis, and Tissue Inhibitor of Metalloproteinases Expression. Am J Resp Cell Mol 24, 591–598 (2001).

57. Weigle, S., Martin, E., Voegtle, A., Wahl, B. & Schuler, M. Primary cell-based phenotypic assays to pharmacologically and genetically study fibrotic diseases in vitro. J Biological Methods 6, e115 (2019).

58. Chantranupong, L. et al. Rapid purification and metabolomic profiling of synaptic vesicles from mammalian brain. Elife 9, e59699 (2020).

59. Love, M. I., Huber, W. & Anders, S. Moderated estimation of fold change and dispersion for RNA-seq data with DESeq2. Genome Biol 15, 550 (2014).

60. Yang, J. et al. The I-TASSER Suite: protein structure and function prediction. Nat Methods 12, 7–8 (2015).

61. Jo, S., Kim, T., Iyer, V. G. & Im, W. CHARMM-GUI: A web-based graphical user interface for CHARMM. J. Comput. Chem. 29, 1859–1865 (2008).

62. Sparks, R. P. et al. An Allosteric Binding Site on Sortilin Regulates the Trafficking of VLDL, PCSK9, and LDLR in Hepatocytes. Biochemistry 59, 4321–4335 (2020).

63. Huang, J. et al. CHARMM36m: an improved force field for folded and intrinsically disordered proteins. Nat Methods 14, 71–73 (2017).

64. Zhang, Y. I-TASSER server for protein 3D structure prediction. Bmc Bioinformatics 9, 40 (2008).

