## Supplemental Material for "Screening the human druggable genome identifies ABHD17B as an anti-fibrotic target in hepatic stellate cells"

Supplementary Materials & Method

*siRNAs information*

The siRNAs targeting *FAM160B2*, *MGAT5B*, *TMEM134*, *TTTY2* and *TTY2B* were provided by the Institute of Chemistry and Cell Biology (ICCB)-Longwood screening facility, and the relevant information is provided in the supplementary tables and in the table below.

| **Gene Symbols** | **Accession Numbers** | **si** | **Sequence** | **Anti-sense Sequence** |
| --- | --- | --- | --- | --- |
| FAM160B2 | NM_022749 | 1 | GGAGACAGGCUAUGACACA | UGUGUCAUAGCCUGUCUCC |
|  |  | 2 | GCAGAGAAUCCAGAGGGUA | UACCCUCUGGAUUCUCUGC |
|  |  | 3 | GGCAUCAGCUGGAGGUUAC | GUAACCUCCAGCUGAUGCC |
|  |  | 4 | CCGAGAUCGUCAACAGUUU | AAACUGUUGACGAUCUCGG |
| MGAT5B | NM_144677 | 1 | GGACGGAACCUGCGUACAA | UUGUACGCAGGUUCCGUCC |
|  |  | 2 | CAAGUUCCCUGACUGCUCA | UGAGCAGUCAGGGAACUUG |
|  |  | 3 | GAACGUCUCCGACAUCGCU | AGCGAUGUCGGAGACGUUC |
|  |  | 4 | GGACAGUCGACUACAACAA | UUGUUGUAGUCGACUGUCC |
| TMEM134 | NM_001078650 | 1 | CAACACCCUUUGAUCCAGA | UCUGGAUCAAAGGGUGUUG |
|  |  | 2 | CCUGGAGUCUAUCACGUGA | UCACGUGAUAGACUCCAGG |
|  |  | 3 | GGUUCGAGGUGGCUGACGA | UCGUCAGCCACCUCGAACC |
|  |  | 4 | AUUAAUUCCACCAGAGGCA | UGCCUCUGGUGGAAUUAAU |
| TTTY2 | NR_001536 | 1 | GGUCAAUAAUUUUACGGUA | UACCGUAAAAUUAUUGACC |
|  |  | 2 | CCUAUUAAAUUUACCUCGA | UCGAGGUAAAUUUAAUAGG |
|  |  | 3 | UGAGAGACCCGUUCGACAU | AUGUCGAACGGGUCUCUCA |
|  |  | 4 | CCCCAGGAAUAAACGGCAA | UUGCCGUUUAUUCCUGGGG |
| TTTY2B | NR_003590 | 1 | GGUCAAUAAUUUUACGGUA | UACCGUAAAAUUAUUGACC |
|  |  | 2 | CCUAUUAAAUUUACCUCGA | UCGAGGUAAAUUUAAUAGG |
|  |  | 3 | UGAGAGACCCGUUCGACAU | AUGUCGAACGGGUCUCUCA |
|  |  | 4 | CCCCAGGAAUAAACGGCAA | UUGCCGUUUAUUCCUGGGG |

The siRNAs used in the validation experiments were purchased from Horizon Discovery as listed below.

| **Full name** | **Abbreviation** | Cat. # |
| --- | --- | --- |
| siGENOME non-targeting control siRNA#1 | NTC si1 | D-001210–01 |
| siGENOME non-targeting control siRNA#2 | NTC si2 | D-001210–02 |
| siGENOME non-targeting control siRNA#3 | NTC si3 | D-001210–03 |
| siGENOME non-targeting control siRNA#4 | NTC si4 | D-001210–04 |
| siGENOME non-targeting control siRNA#5 | NTC si5 | D-001210–05 |
| siGENOME human *ACTA2* siRNA 1 | *ACTA2* si1 | D-003450-01 |
| siGENOME human *ACTA2* siRNA 2 | *ACTA2* si2 | D-003450-02 |
| siGENOME human *ASAH1* siRNA 1 | *ASAH1* si1 | D-005228-01 |
| siGENOME human *ASAH1* siRNA 4 | *ASAH1* si4 | D-005228-04 |
| siGENOME human *PSMC4* siRNA 1 | *PSMC4* si1 | D-009261-17 |
| siGENOME human *PSMC4* siRNA 2 | *PSMC4* si4 | D-009261-02 |
| siGENOME human *PLK1* SMARTpool | *PLK1* siP | M-003290-01 |
| siGENOME human *KIF11* SMARTpool | *KIF11* siP | M-003317-01 |
| siGLO RISC-free siRNA | siGLO | D-001600-01 |
| siGENOME human *GAPDH* SMARTpool | *GAPDH* siP | M-004253-02 |
| siGENOME human *NFkB1* SMARTpool | *NFkB1* siP | M-003520-01 |
| siGENOME human *NFkB2* SMARTpool | *NFkB2* siP | M-003918-02 |
| siGENOME human *NR1H2* SMARTpool | *NR1H2* siP | M-003412-03 |
| siGENOME human *NR1H3* SMARTpool | *NR1H3* siP | M-003413-01 |
| siGENOME human *UBB* SMARTpool | *UBB* siP | M-013382-01 |
| siGENOME human *TLR6* SMARTpool | *TLR6* siP | M-005156-01 |
| siGENOME human *AURKC* SMARTpool | *AURKC* siP | M-019573-04 |
| siGENOME human *AKAP11* SMARTpool | *AKAP11* siP | M-009277-01 |
| siGENOME human *ABHD17B* siRNA#1 | *ABHD17B* si1 | D-005809-01 |
| siGENOME human *ABHD17B* siRNA#4 | *ABHD17B* si4 | D-005809-04 |
| siGENOME human *ABHD17A* SMARTpool | *ABHD17A* siP | M-005947-02 |
| siGENOME human *ABHD17C* SMARTpool | *ABHD17C* siP | M-005929-02 |
| siGENOME human *MYO1B* SMARTpool | *MYO1B* siP | M-023110-00 |
| ON-TARGETplus human ABHD17B pool | ABHD17B siP | L-005809-01 |
| ON-TARGETplus non-targeting control pool | NTC siP | D-001810-10 |

*Lipid accumulation assay based on Hoechst and Bodipy staining*

HSCs transfected with siRNAs or treated with compounds were fixed with 4% paraformaldehyde (Electron Microscopy Sciences, Cat. # 15710) for 15 min at room temperature (RT). Cells were washed with DPBS and stained with Bodipy 493/503 (0.25 μg/mL, Invitrogen, Cat. # D3922) and Hoechst (5 μg/ mL, Invitrogen, Cat. # H1399) for 45 min at RT. Images were taken with the ImageXpress Micro Confocal (Molecular Devices) at the Institute of Chemistry and Cell Biology (ICCB)-Longwood screening facility and analyzed using the MetaXpress software. Cells with a cytoplasmic Bodipy staining intensity higher than the cutoff were defined as Bodipy-positive cells. Cells were counted and normalized to the total nuclei count in the same microscopic field to calculate the percentage of Bodipy positive cells. The cutoff was adjusted for each plate, so that there are about 10-20% Bodipy-positive cells in the DMSO condition and 80-90% Bodipy-positive cells in the nortriptyline-treated positive control wells. For the primary screen, the results were scanned for outliers, which were then corrected based on the method described in the Supplementary Materials & Methods. The score was then calculated based on the averaged percent positive cells for the siRNA compared to the baseline of the plate, as well as the correlation among three replicates. Toxicity was calculated based on total cell numbers.

*Outlier correction in the primary and validation screens*

To determine if the replicates of a specific library well contain outliers, the interquartile range (IQR) was calculated using all the replicates for each siRNA. The minimum IQR that was considered an outlier among all IQRs was used to set up the threshold to correct for outliers. In the primary screen, siRNAs that had an IQR larger than 11.3891 were corrected. To correct an outlier, the Tukey’s method was applied, which uses a cleaning parameter to detect outliers. The cleaning parameter is 1.5 x 11.3891, following the same logic as the outlier detection when plotting a boxplot. If any value was larger than the median +/- the cleaning parameter, that value was set to be exactly the median +/- the cleaning parameters. This method allows us to keep some of the variability while reducing the extreme variance that an outlier can generate.

*High-throughput evaluation of mRNA expression and knockdown efficiency*

HSCs were transfected with siRNAs as described above. 72 hr after transfection, cells were harvested using the Cells-to-CT 1-Step Taqman Kit (Invitrogen, Cat. # A25603) as previously described^1^ except that lysis was performed with 15 µL lysis buffer (plus DNase) for 5 min at RT, and then the reaction was stopped by adding 1.5 µL stop solution and incubating for 2 min at RT. To quantify the depletion of each target gene, customized 384-well plates arrayed with Taqman probes for each mRNA target gene were purchased from Thermo Fisher Scientific. Reverse transcription and amplification of cDNA was performed with Taqman 1-Step qRT-PCR mix supplied with the kit. *ACTA2*, *COL1A1* and target mRNA were each quantified in the same well with the endogenous control *PSMB2*. *ACTA2*, *COL1A1*, and target mRNA Taqman probes are FAM-labeled, while the *PSMB2* probe is VIC-labeled and primer-limited.

The mRNA expression results for *ACTA2* (normalized to *PSMB2*) and *COL1A1* (normalized to *PSMB2*) were analyzed using two methods: the linear regression strategy and the standard ΔΔCt method. For the linear regression strategy, the data was fit to a linear model: Ct_*ACTA2* (or *COL1A1*) ~ Ct_*PSMB2* + plate + siRNA. The estimate for each experimental siRNA was compared to that of the control siRNA to calculate the relative fold change. The knockdown efficiency for each target gene was calculated using the standard ΔΔCt method with *PSMB2* as the endogenous control.

*TaqMan Real-time PCR Assays for analyzing mRNA levels*

|  | **Assay ID** |
| --- | --- |
| Human *ACTA2* | Hs00426835_g1 |
| Human *COL1A1* | Hs00164004_m1 |
| Human *PSMB2* | Hs01002946_m1 |
| Human *ASAH1* | Hs00602774_m1 |
| Human *PLK1* | Hs00983227_m1 |
| Human *UBB* | Hs00430290_m1 |
| Human *ABHD17B* | Hs00925211_m1 |
| Human *GAPDH* | Hs02786624_g1 |
| Human *COL3A1* | Hs00943809_m1 |
| Human *ABHD17A* | Hs07290154_g1 |
| Human *ABHD17C* | Hs01593305_m1 |
| Human *AURKC* | Hs00152930_m1 |
| Human *AKAP11* | Hs01568654_m1 |
| Human *TLR6* | Hs04975840_m1 |
| Human *NFkB1* | Hs00765730_m1 |
| Human *NFkB2* | Hs01028890_g1 |
| Human *NR1H2* | Hs01027208_m1 |
| Human *NR1H3* | Hs00172885_m1 |
| Human *POLR2A* | Hs00172187_m1 |
| Human *MYO1B* | Hs00362654_m1 |
| Human *PSMC4* | Hs00197826_m1 |
| Human *TIMP1* | Hs00171588_m1 |
| Human *TGFBR1* | Hs00610320_m1 |
| Mouse *Gapdh* | Mm99999915_g1 |
| Mouse *Col1a1* | Mm00801666_g1 |
| Mouse *Acta2* | Mm00725412_s1 |
| Mouse *Timp1* | Mm01341361_m1 |
| Mouse *Tgfb1* | Mm01178820_m1 |
| Mouse *Il1b* | Mm00434228_m1 |
| Mouse *Abhd17b* | Mm01197077_m1 |

*Analysis of mice single-cell RNA-sequencing data*

To delineate mouse *Abhd17b* expression across various cell types, single-cell RNA-sequencing data from oil-treated control and CCl4-treated mice were obtained from the NCBI GEO database (GSE171904 dataset)^2^. Cell Ranger (version 7.0.1; 10x Genomics) was used to process the raw sequencing data with default parameters, generating expression matrices using the cellranger count pipeline. These matrices from the two conditions were then aggregated with the cellranger aggr pipeline. The aggregated data was subsequently imported into R and analyzed with the Seurat package (version 5.0.0) to perform quality control, normalization, and downstream analyses. Cell viability was ensured by retaining only cells with more than 200 expressed genes, and only genes present in at least three cells were included. Data were log-normalized, and cell type relationships and UMAP coordinates were derived from the RDS files accompanying the GSE171904 dataset. Dot plots were generated by the "scanpy" Python package, and gene expression levels in dot plots represent mean expression across all cells in a cluster.

*Generation of ABHD17B S170A mutant*

DNA sequences that encode the wildtype amino acid sequence of human *ABHD17B* and encode a mutation of Ser 170 to Ala in *ABDH17B* were synthesized by GENScript. The cDNAs also contain additional identical nucleotide modifications that do not affect amino acid sequence (**Supplementary Fig. 15**).

*RNA sequencing*

HSCs from donor 3 were transfected with NTC si5, *ABHD17B* si1, and *ABHD17B* si4 in triplicate. Cells were harvested in Trizol reagent, and RNA was purified through a Direct-zol RNA Miniprep Kit (Cat. #R2050). All samples had RIN scores greater than 9 (Agilent 4150 TapeStation System, G2992 AA) and underwent PolyA-selection and stranded library preparation prior to sequencing at 150 paired end reads (Genewiz from Azenta Life Sciences). RNA-seq counts were generated by bcbio-nextgen using salmon^3^. Counts were imported into R using tximport^4^ and DESeq2^5^. Gene annotations for pseudo-alignment were obtained from Ensembl, version Homo_sapiens.GRCh38.98. AnnotationHub was used to obtain annotations for the Rscripts. Data manipulation and plots were done using Tidyverse^6^. Differential expression analysis was performed using DESeq2^5^ with lfcShrinkage using [apeglm]^7^. Genes were defined as repressed by the following criteria: Adjusted p value <0.05 for both si1 and si4 and (log2FC-siRNA1_vs_ctrl <= -0.585 AND log2FC-siRNA4_vs_ctrl <= 0) or (log2FC-siRNA1_vs_ctrl <= 0 AND log2FC-siRNA4_vs_ctrl <= -0.585). Heatmaps were generated by pheatmap^8^. PCA plots were generated using custom code. For functional analysis clusterProfiler^9^ with GOSemSim^10,11^ and DOSE^12^ were used.

*Hepatic hydroxyproline*

To quantify collagen level, mouse liver samples were isolation from the same region of the left liver lobe^13^. Isolated samples were homogenized and processed to evaluate hydroxyproline concentration using hydroxyproline assay kits (Sigma-Aldrich, Cat. # MAK008) following manufacturer’s instruction.

*Collagen proportionate area (CPA)* *and immunohistochemistry (IHC)*

Livers were fixed in 4% paraformaldehyde at 4° C. After dehydration through graded ethanol and paraffin embedment, samples were sectioned at 5 μm. CPA was measured as described previously^14^. Pico-Sirius red staining was performed using the left liver lobe from formalin fixed paraffin embedded sections (iHisto). Whole sections were scanned and loaded into ImageJ to calculate the ratio of collagen positive area against the total parenchyma area and expressed as a percentage. Immunohistochemistry (IHC) was conducted on paraffin sectioned samples following standard IHC procedure with antigen recovery. The following antibodies were used: PDGF Receptor β (28E1) antibody (Cell Signaling Technology, Cat. # 3169), α-Smooth Muscle Actin (D4K9N) antibody (Cell Signaling Technology, Cat. # 19245), and Cytokeratin 19 antibody [EP1580Y] (Abcam, Cat. # ab52625).

*Statistical analysis and figure preparation*

Data involving comparisons of more than two groups were analyzed in GraphPad Prism using one-way ANOVA test, except for the scar-in-a-jar assay, which were analyzed using Dunnetts multiple comparison test. Mouse experiments and qRT-PCR results comparing two groups of data were analyzed by upaired two-tailed student’s t-test. In all figures where applicable, ns indicates not significant, * indicates p<0.05, ** indicates p<0.01, *** indicates p<0.001, and **** indicates p<0.0001. In all bar graphs, each dot represents one biological replicate. Error bars represent mean ± SEM, except for the scar-in-a-jar assay and the dose response curve in Fig. 3C, where error bars represent mean ± SD. Results are representative of 2-3 independent experiments.

Supplementary Figures

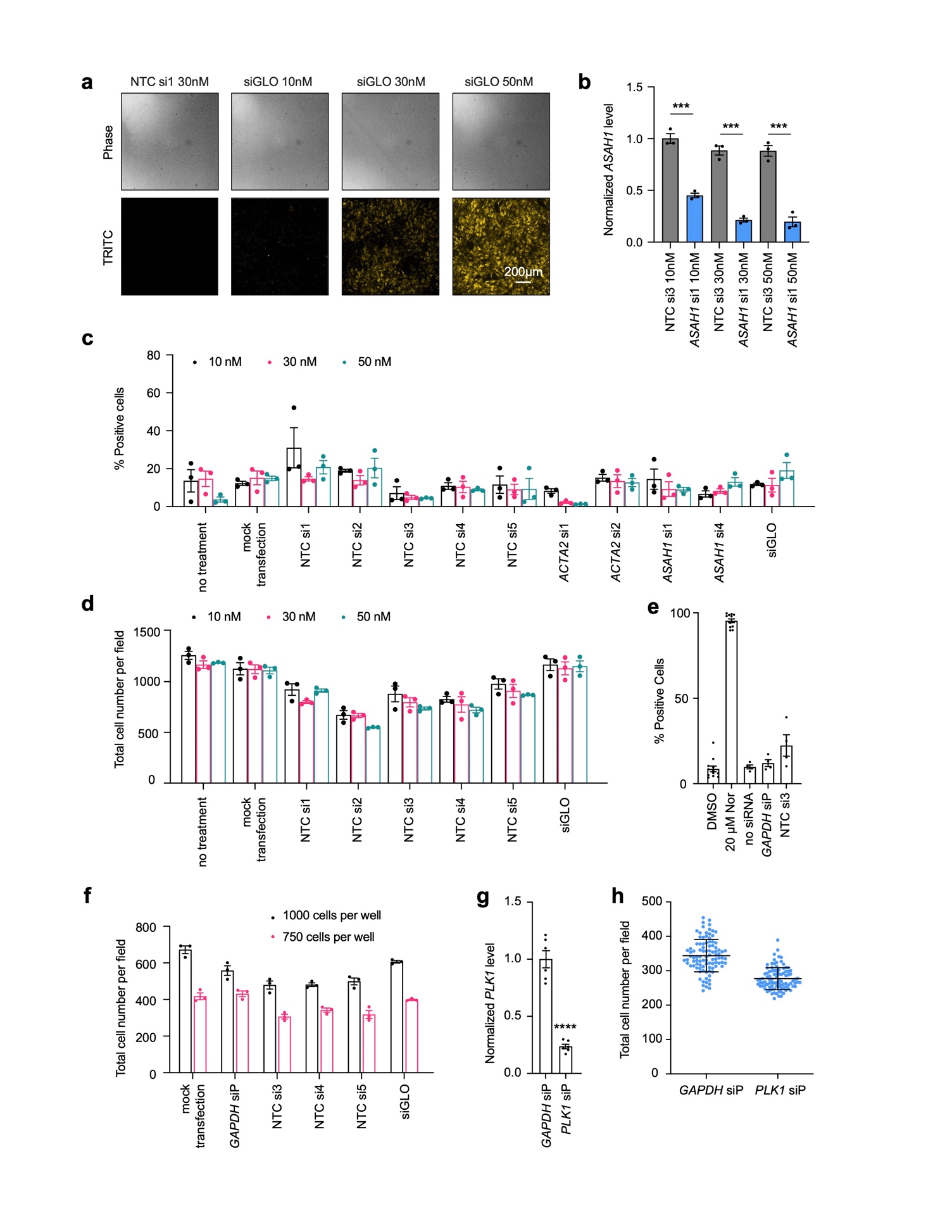

**Supplementary Fig. 1. Optimization of siRNA transfection protocol in human primary HSCs.** (a) Primary HSCs from donor 1 were transfected with non-targeting (NTC) siRNA or siGLO at the indicated concentrations. Cells were imaged 72 hr after transfection. Representative images are shown. Scale bar represents 200 µm. (b) Primary HSCs from donor 1 were transfected with non-targeting siRNA or siRNA targeting *ASAH1*^15^ at indicated concentrations, and cells were lysed for RNA extraction 72 hr after transfection. The knockdown efficiency was determined by qRT-PCR. Each dot represents one biological replicate. Error bars represent mean ± SEM. *** indicates p<0.001 (n=3, unpaired two-tailed student’s t-test). (c-d) Primary HSCs from donor 1 were transfected with different non-targeting control (NTC) siRNAs, siRNAs targeting *ACTA2*, *ASAH1*, or siGLO at indicated concentrations, and cells were fixed for Hoechst and Bodipy staining 72 hr after transfection. Cells were not treated with transfection reagents nor siRNAs in the “no treatment” group, while cells were treated with transfection reagents without siRNAs in the “mock transfection” group. Each dot represents one biological replicate (n=3). Error bars represent mean ± SEM. c: percentage of Bodipy positive cells. d: total count of nuclei per microscopic imaging field. (e) Primary HSCs from donor 1 were treated with DMSO or nortriptyline for 48 hr (used as controls for Bodipy staining) or transfected with siRNAs as indicated for 72 hr. Cells were then fixed and stained with Hoechst and Bodipy to determine the percentage of Bodipy positive cells. Each dot represents one biological replicate. Error bars represent mean ± SEM. (f) Primary HSCs were seeded at two densities in 384-well plates and transfected with 25 nM siRNAs as indicated, and cells were fixed for Hoechst staining 72 hr after transfection. Each dot represents one biological replicate (n=3). Error bars represent mean ± SEM. (g-h) Primary HSCs were transfected with 25 nM pooled siRNAs targeting *PLK1*, and cells were lysed for RNA extraction (g) or fixed for Hoechst staining (h) 72 hr after transfection. g: The knockdown efficiency was determined by qRT-PCR. Each dot represents one biological replicate. Error bars represent mean ± SEM. **** indicates p<0.0001 (n=6, t test). h: The number of cells was determined by nuclei count. Each dot represents one biological replicate (n=96). Error bars represent mean ± SD.

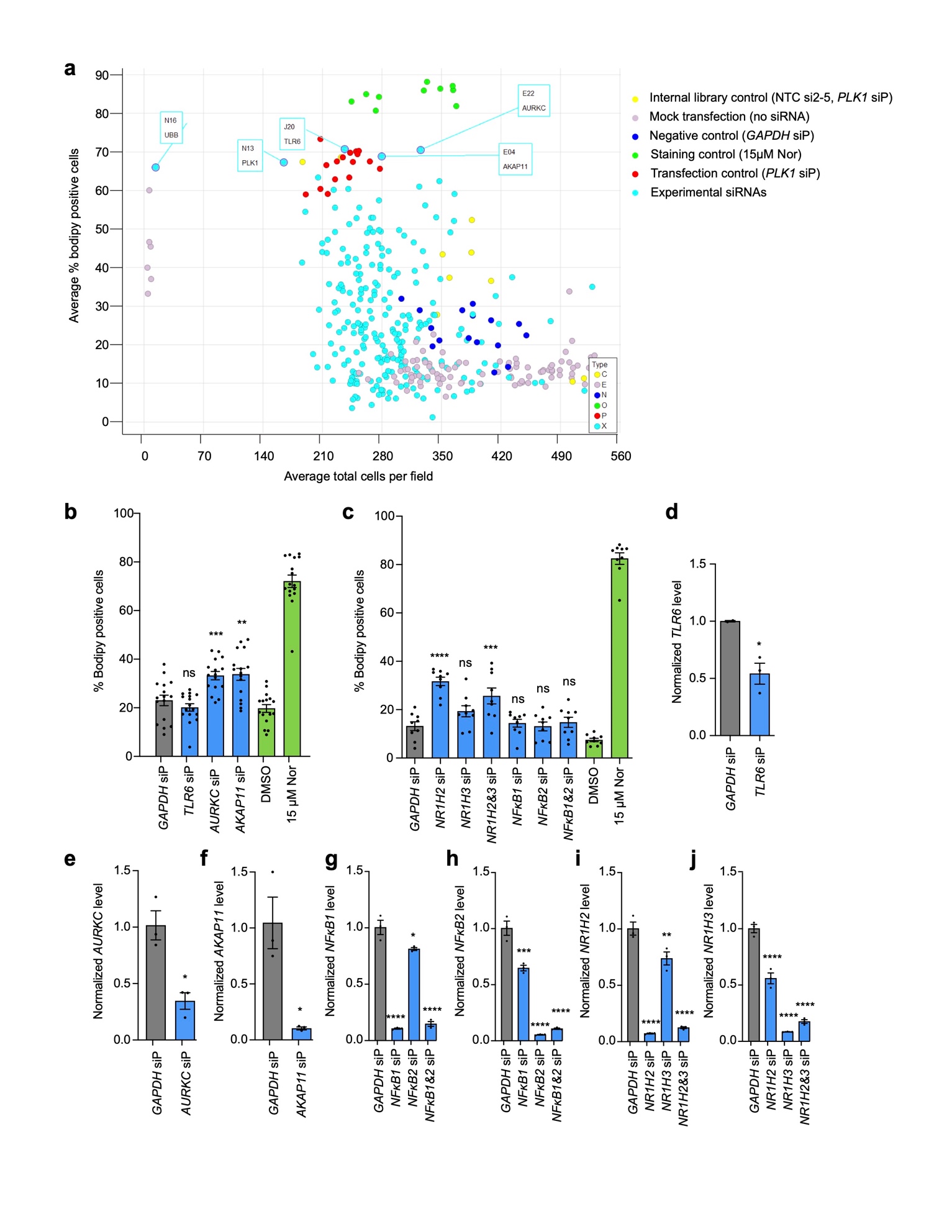

**Supplementary Fig. 2. Pilot siRNA library screen and evaluation of potential positive controls.** (a) Summary of the pilot siRNA screen results. Each dot represents the average % Bodipy positive cells and average total cells per field of three replicates of an experimental or control well. All wells from the pilot library plate are shown. The experimental siRNAs with high % Bodipy positive cells or low cell numbers are labeled with well ID and target gene symbol. (b-c) HSCs were treated with siRNAs, DMSO, or nortriptyline (Nor) as indicated, and 72 hr later, cells were fixed and stained with Hoechst and Bodipy. Each dot represents one biological replicate. Error bars represent mean ± SEM. ns indicates not significant, ** indicates p<0.01, *** indicates p<0.001, and **** indicates p<0.0001 compared to the negative control *GAPDH* siP (n=16, one-way ANOVA test). (d-j) HSCs were treated with siRNAs, DMSO, or nortriptyline as indicated, and 72 hr later, cells were lysed for RNA extraction. Knockdown efficiency was evaluated by qRT-PCR. Each dot represents one biological replicate. Error bars represent mean ± SEM. * indicates p<0.05, ** indicates p<0.01, *** indicates p<0.001, and **** indicates p<0.0001 compared to the negative control *GAPDH* siP (n=3, t test for d-f and one-way ANOVA test for g-j).

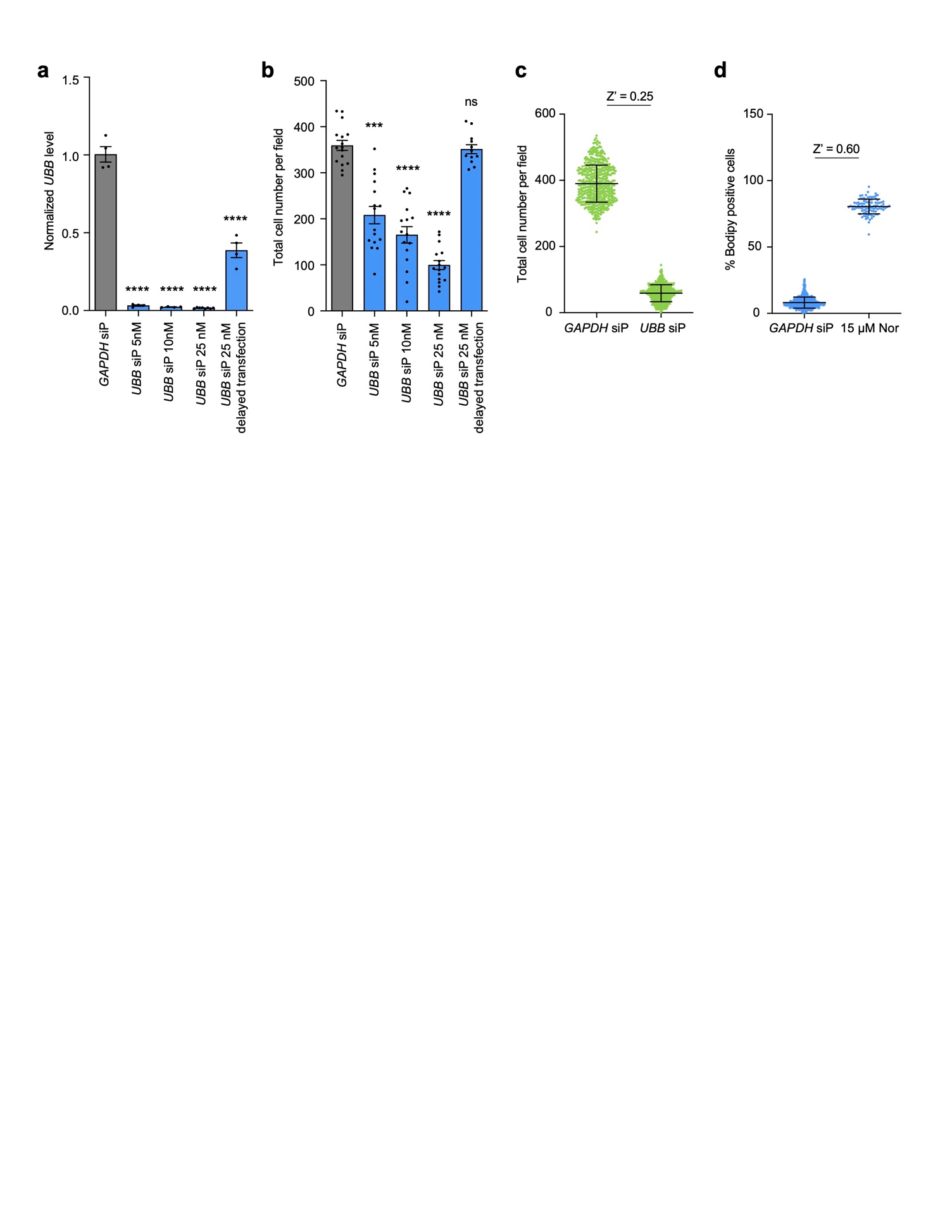

**Supplementary Fig. 3. *UBB* depletion as an indicator of transfection efficiency.** (a-b) Primary human HSCs from donor 1 were transfected with pooled *UBB* siRNAs (siP) at the concentrations indicated. For the “delayed transfection” test condition, transfection reagents were added later than the optimal time frame to confirm that the reagents were not toxic by themselves. Cells were lysed for RNA extraction (a) or fixed and stained with Hoechst (b) 72 hr after transfection. Each dot represents one biological replicate. Error bars represent mean ± SEM. ns indicates not significant, *** indicates p<0.001, and **** indicates p<0.0001 compared to the negative control *GAPDH* siP (n≥4 as indicated by the number of dots, one-way ANOVA test). a: depletion efficiency was determined by qRT-PCR. b: Total cell number per field was quantified based on nuclei count. (c) Primary human HSCs from donor 1 were transfected with pooled *GAPDH* or *UBB* siRNAs at 25 nM. Cells were fixed and stained with Hoechst 72 hr after transfection to quantify cell number. Each dot represents one biological replicate (n=528). Error bars represent mean ± SD. (d) Primary human HSCs from donor 1 were treated with pooled *GAPDH* siRNAs at 25 nM or nortriptyline (Nor) at 15 µM for 72 hr before cells were fixed and stained with Bodipy and Hoechst. Each dot represents one biological replicate (n=528 for *GAPDH* siP, n=96 for Nor). Error bars represent mean ± SD.

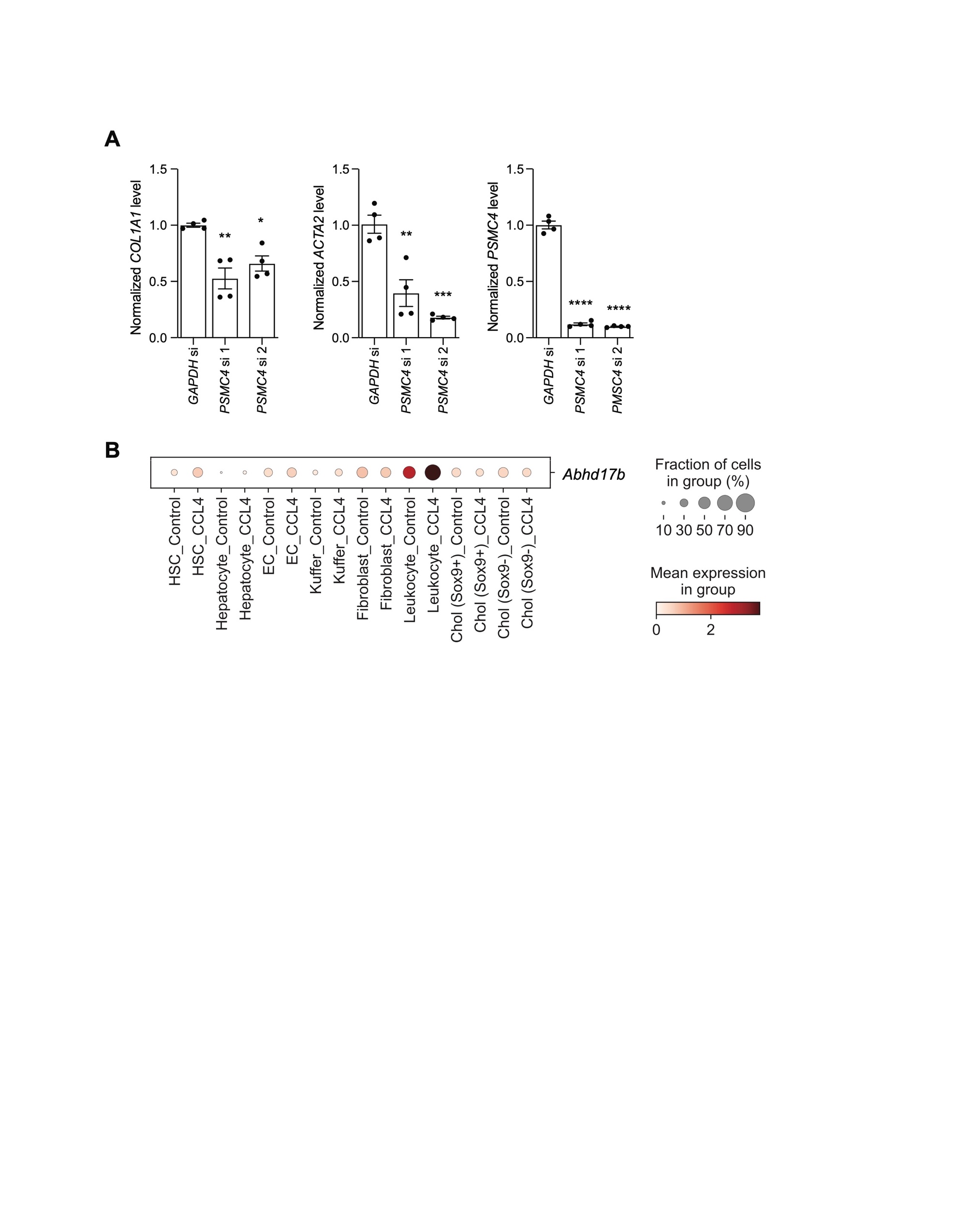

**Supplementary Fig. 4. Depletion of *PSMC4* reduces expression of *COL1A1* and *ACTA2*.** Primary human HSCs from donor 1 were treated with pooled siRNAs targeting *GAPDH* and individual siRNAs targeting *PSMC4* (si 1 and si 2) at 25 nM for 72 hr before analysis of expression for the indicated genes by qRT-PCR. Each dot represents one well. * indicates p <0.05, ** indicates p<0.01, *** indicates p<0.001, and **** indicates p <0.0001 (one-way ANOVA). Error bars represent mean ± SEM.

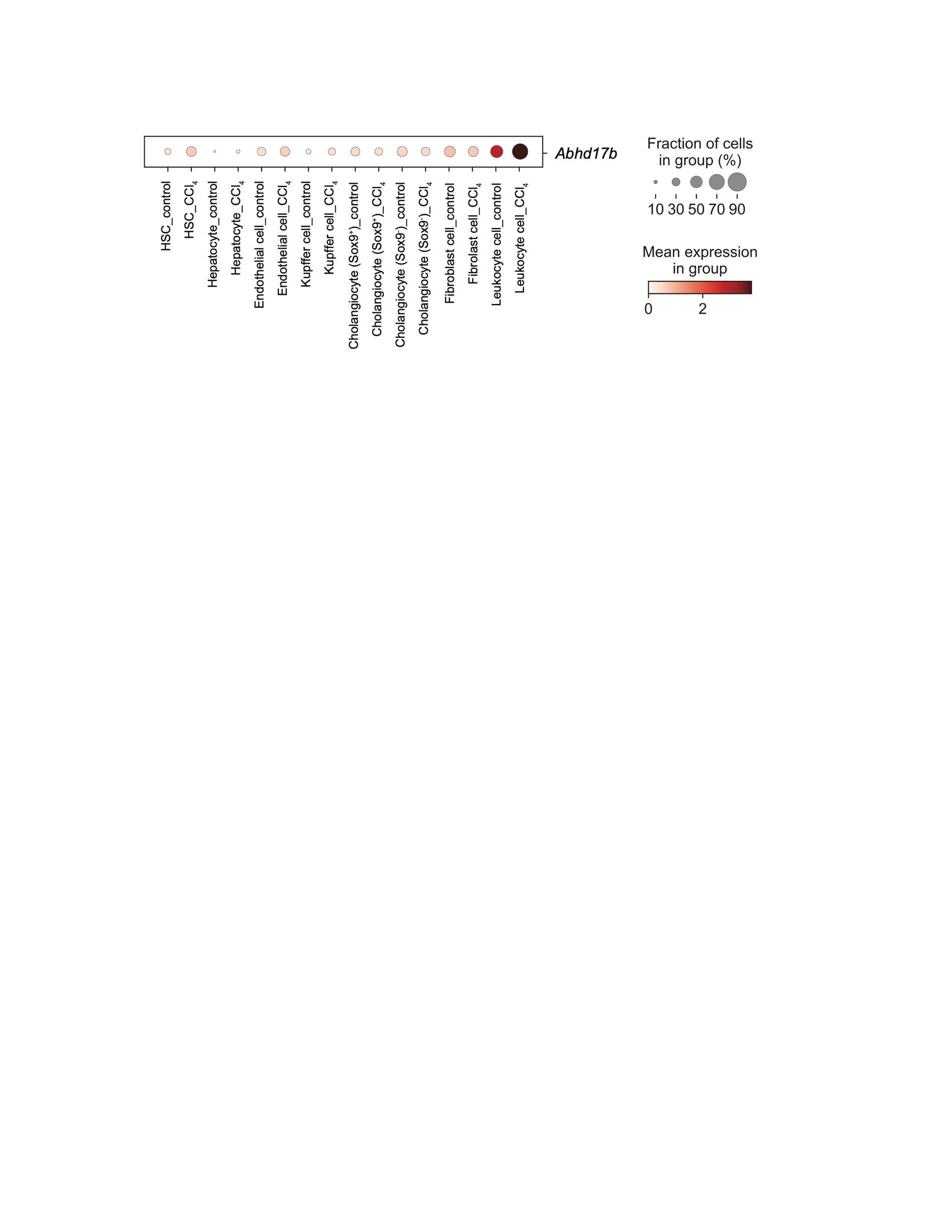

**Supplementary Fig. 5. Expression of *Abhd17b* in different cell types from the livers of CCl_4_-treated and control mice.** Single-cell RNA sequencing results^2^ were analyzed as described in supplementary methods. Circle size represents the fraction of cells expressing *Abhd17b*, and color indicates mean expression level.

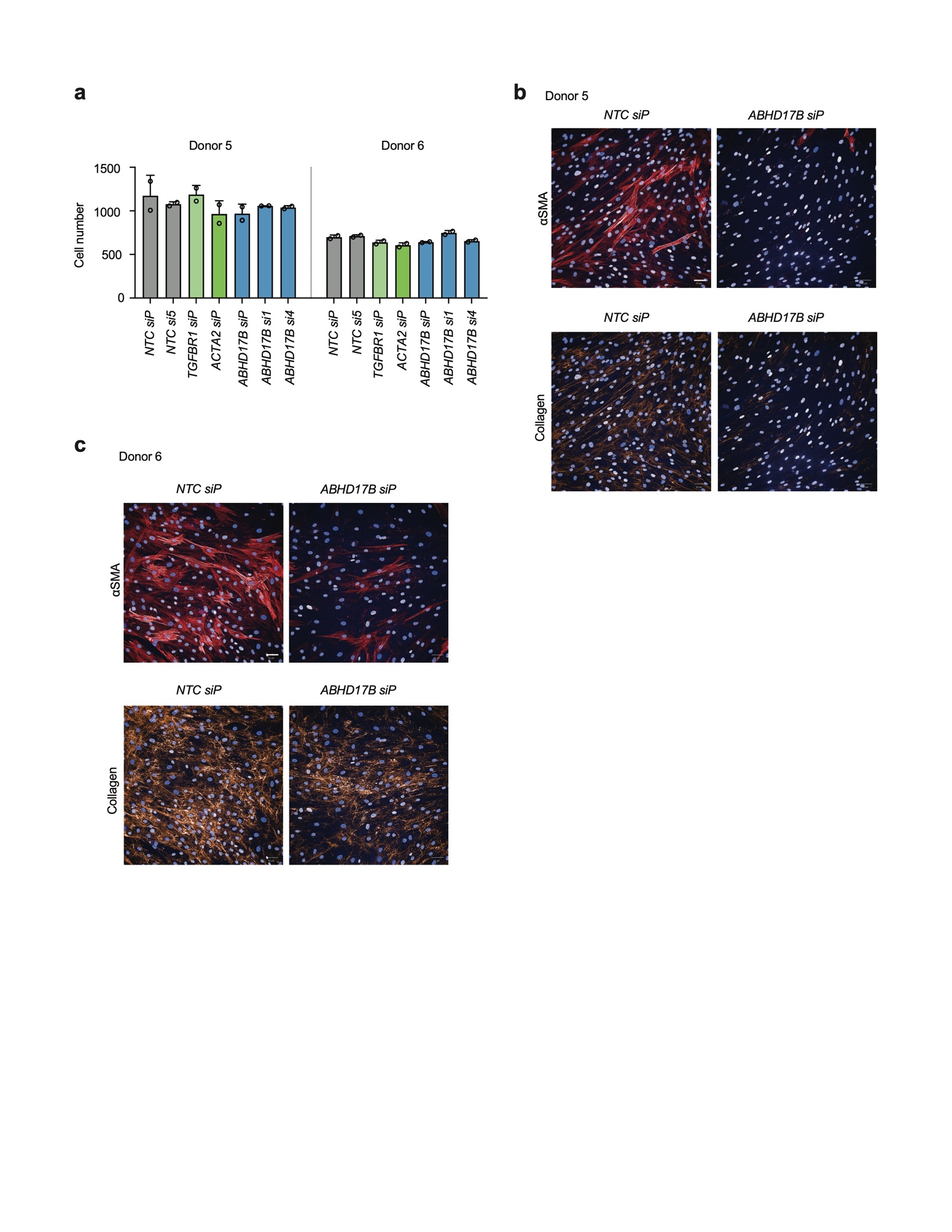

**Supplementary Fig. 6. Scar-in-a-jar assay.** (a) Primary human HSCs from donors 5 and 6 were transfected with siRNAs as indicated and cell numbers were quantified by nuclei (Hoescht) 3 days after transfection. Nontargeting control pooled siRNA (NTC siP) and NTC si5 are compared to pooled *ABHD17B* (siP) siRNA, *ABHD17B si1*, *ABHD17B si4*, and control siRNAs targeting *ACTA2* and *TGFBR1*. Error bars represent mean ± SD for two biological replicates (open circles). (b) HSCs from donor 5 were transfected with pooled nontargeting control (*NTC siP*) and pooled siRNAs targeting *ABHD17B* (*ABHD17B siP*). HSCs were serum starved the day after transfection followed by stimulation with TGF-β in conditions of molecular crowding for 72 hr. αSMA (red) and collagen (orange) were visualized by immunofluorescence in the same field. Nuclei were stained with Hoechst (blue). (c) Analysis was performed for donor 6 as in (b). White bars indicate 50 µm.

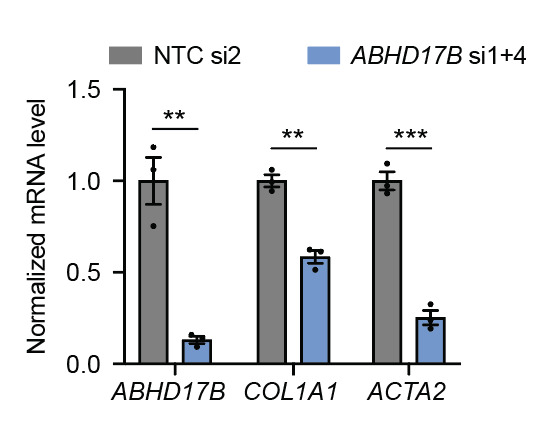

**Supplementary Fig. 7. *ABHD17B* regulates *COL1A1* in lung fibroblasts.** Primary human lung fibroblasts were transfected with siRNAs, serum starved for 24 hr, and then treated with TGF-β for 24 hr. mRNA expression was quantified 72 hr after transfection. Each dot represents one biological replicate. Error bars represent mean ± SEM. ** indicates p<0.01, and *** indicates p<0.001 (n=3, unpaired two-talied student’s t test). Data are representative of three independent experiments. NTC si 2 was used as negative control and compared to depletion of *ABDH17B* with si1 and si4.

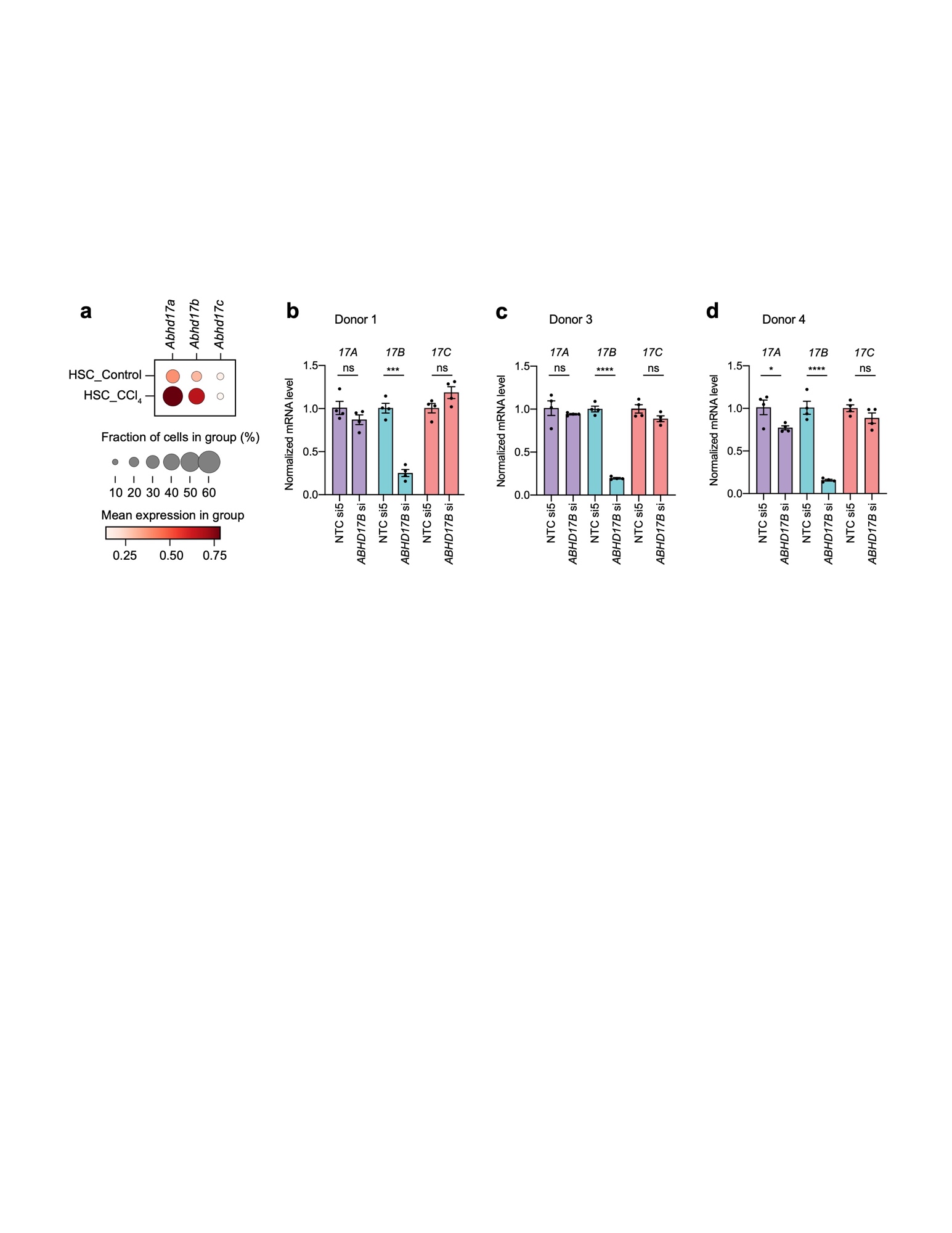

**Supplementary Fig. 8. Abhd17 family gene expression and efficiency of siRNAs targeting each of the ABHD17 family members in HSCs.** (a) Dot plot showing expression of each Abhd17 family gene in HSCs from the livers of CCl_4_-treated and control mice based on single-cell RNA sequencing data^2^. Circle size represents the fraction of cells expressing a gene, and color indicates mean expression level. *ABHD17B* was depleted using pooled siRNAs in HSCs from human donors 1 (b), 3 (c), and 4 (d). *ABHD17A*, *ABHD17B*, and *ABHD17C* levels 72 hr post transfection were quantified by qRT-PCR. Each dot represents one biological replicate. Error bars represent mean ± SEM (n=4). ns indicates not significant, * indicates p<0.05, *** indicates p<0.001, and **** indicates p<0.0001 (one-way ANOVA test).

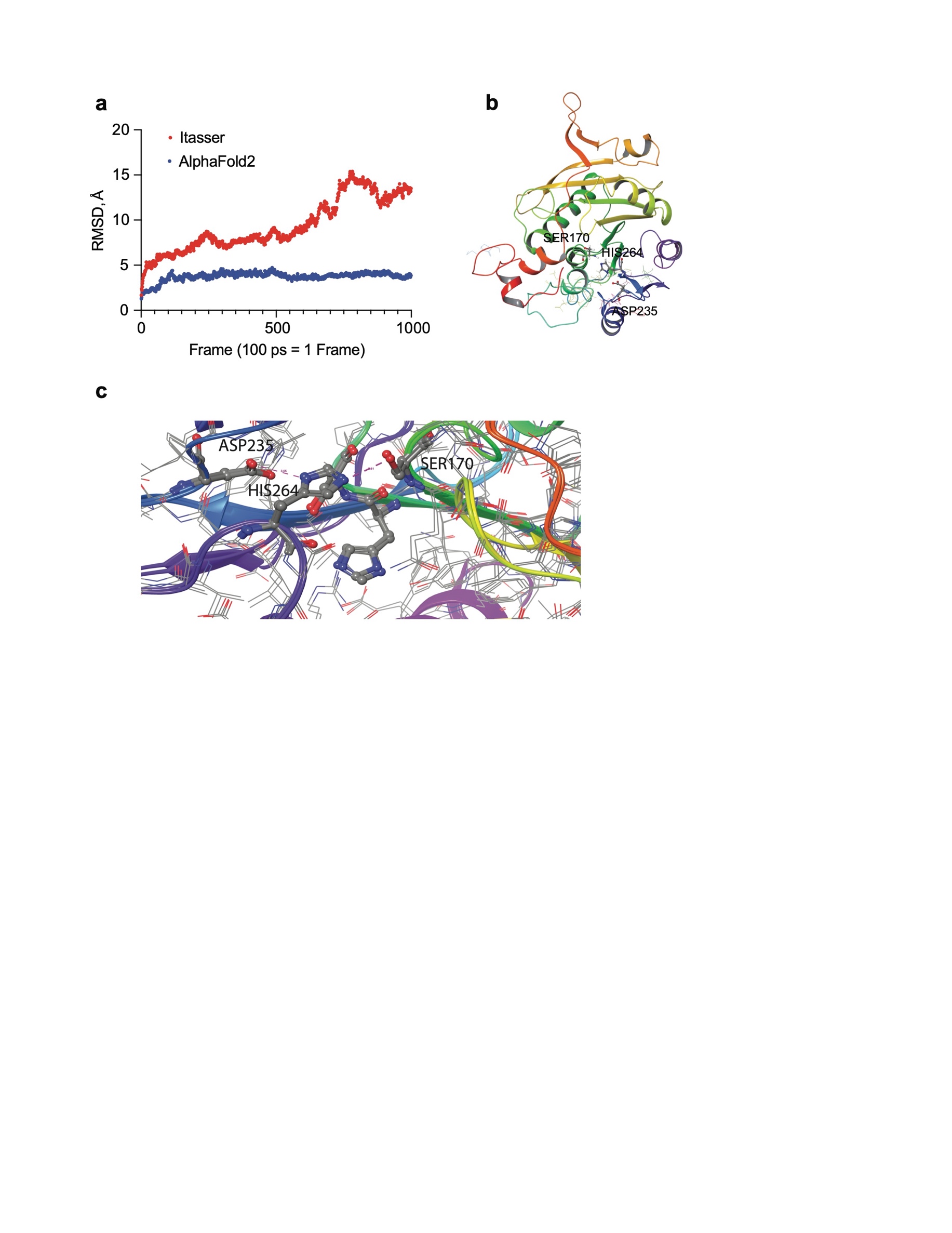

**Supplementary Fig. 9. Modeling of ABHD17B and prediction of Ser170 as required for serine protease activity.** (a) Root mean squared deviation (RMSD) of two simulations of ABHD17B generated through *de novo* protein folding either from I-Tasser server^16^ (red) or AlphaFold2 (blue)^17^. (b) Wildtype AlphaFold2 ABHD17B with proposed catalytic triad (Ser170, His264, and Asp235) indicated. (c) Representation of the potential catalytic triad of ABHD17B from AlphaFold2 minimized with OPLS3^18^ as implemented in Maestro and equilibrated without production step from Namd2, with distance between Asp235 carboxylate oxygens and hydrogen of His264 at 3.05 Angstroms (dashed purple line) as well as distance of deprotonated nitrogen of His264 and hydrogen of hydroxyl of Ser170 at 2.81 Angstroms. The model of ABHD17B is overlayed on a model of ABHD17A showing similarity of structure.

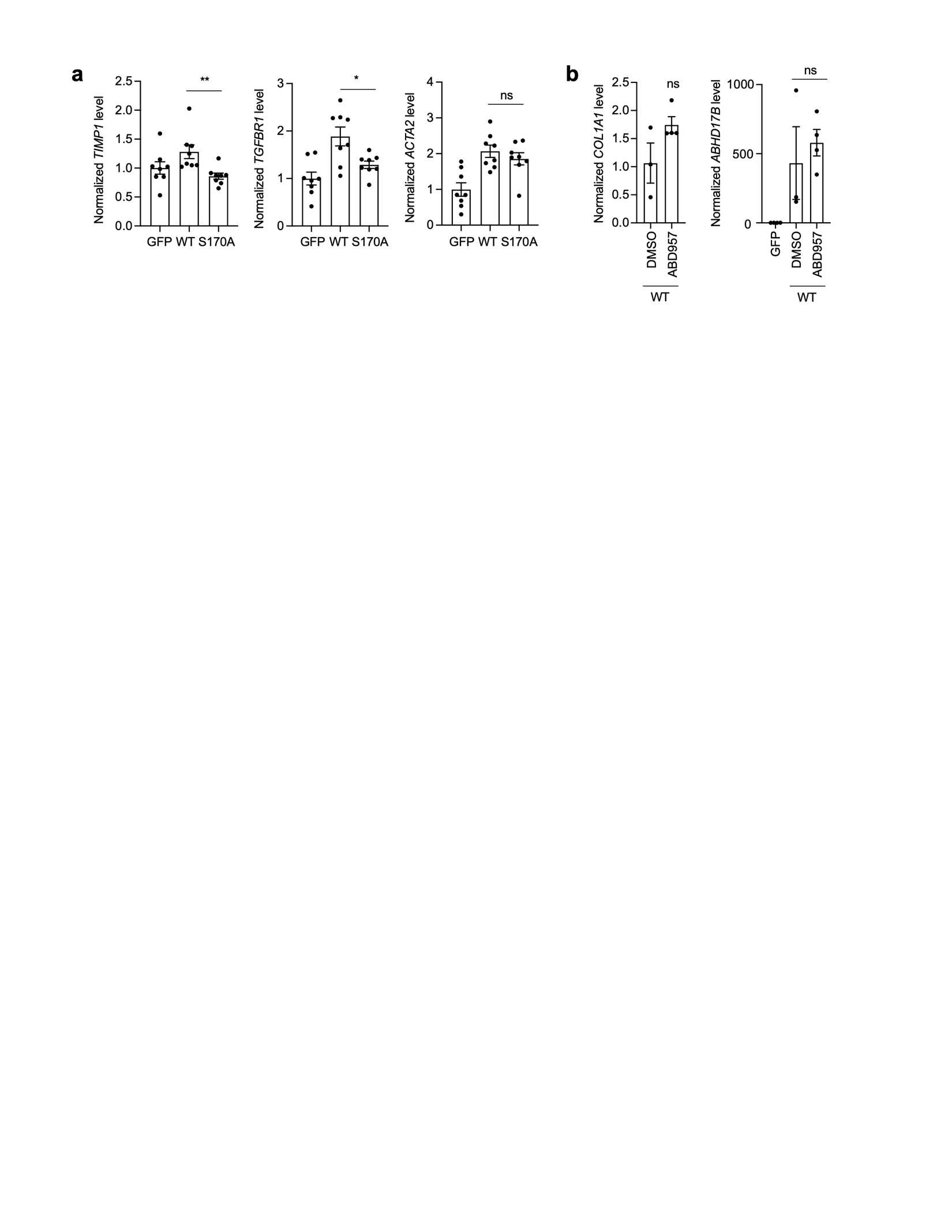

**Supplementary Fig. 10. Effect of ABHD17B overexpression on fibrotic markers and effects of ABD957 on *COL1A1* level in HSCs with overexpression of ABHD17B-WT.** (a) *TIMP1, TGFBR1,* and *ACTA2* were quantified by qRT-PCR in HSCs transduced with lentivirus expressing GFP, ABHD17B-WT or ABHD17B-S170A. Expression was normalized to *PSMB2*. Error bars represent mean ± SEM (n=8). ns indicates not significant, * indicates p<0.05 and ** indicates p<0.01 (2-tailed unpaired t-test between WT and S170A). (b) *COL1A1* and *ABHD17B* were quantified by qRT-PCR in HSCs transduced with lentivirus expressing ABHD17B-WT and treated with DMSO (control) or ABD957 (1 µM). Expression was normalized to *PSMB2*. HSCs transduced with GFP (control) lentivirus was included for reference when quantifying *ABDH17B*. Error bars represent mean ± SEM (n=3 for the DMSO group and n=4 for the other groups). ns indicates not significant (2-tailed unpaired t-test).

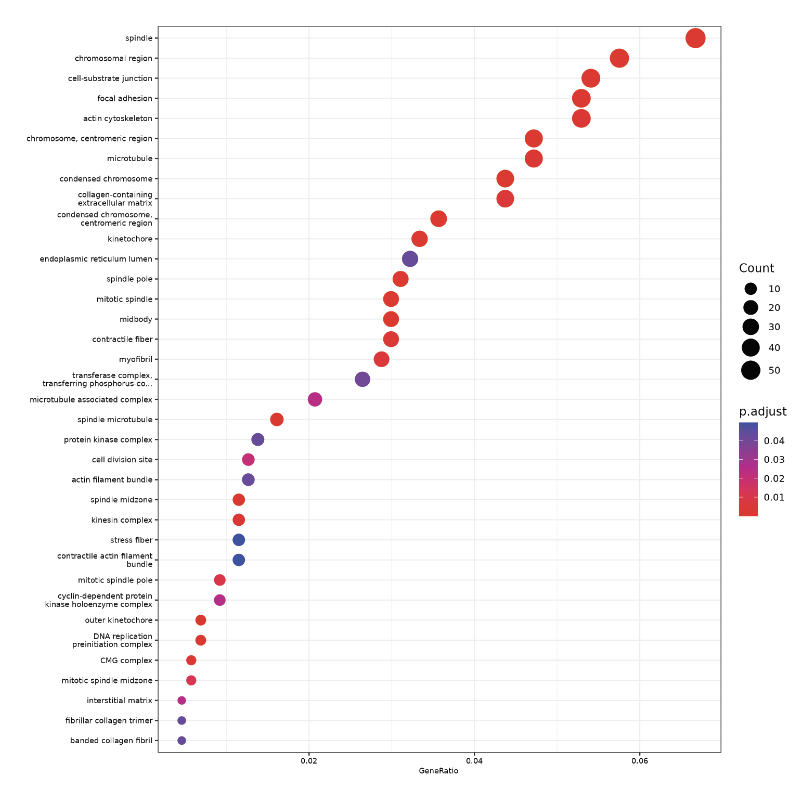

**Supplementary Fig. 11. GO analysis for genes repressed with depletion of *ABHD17B***, **full map from Fig. 5A.** RNA sequencing and differential expression analysis were performed using nontargeting control (NTC) si5, and two siRNAs (siRNA#1 and #4) targeting *ABHD17B*. Dot plot displays the Gene Ontology (GO) terms most enriched following *ABHD17B* depletion. The color of each dot represents the adjusted p-value, and the size of the dot represents gene count.

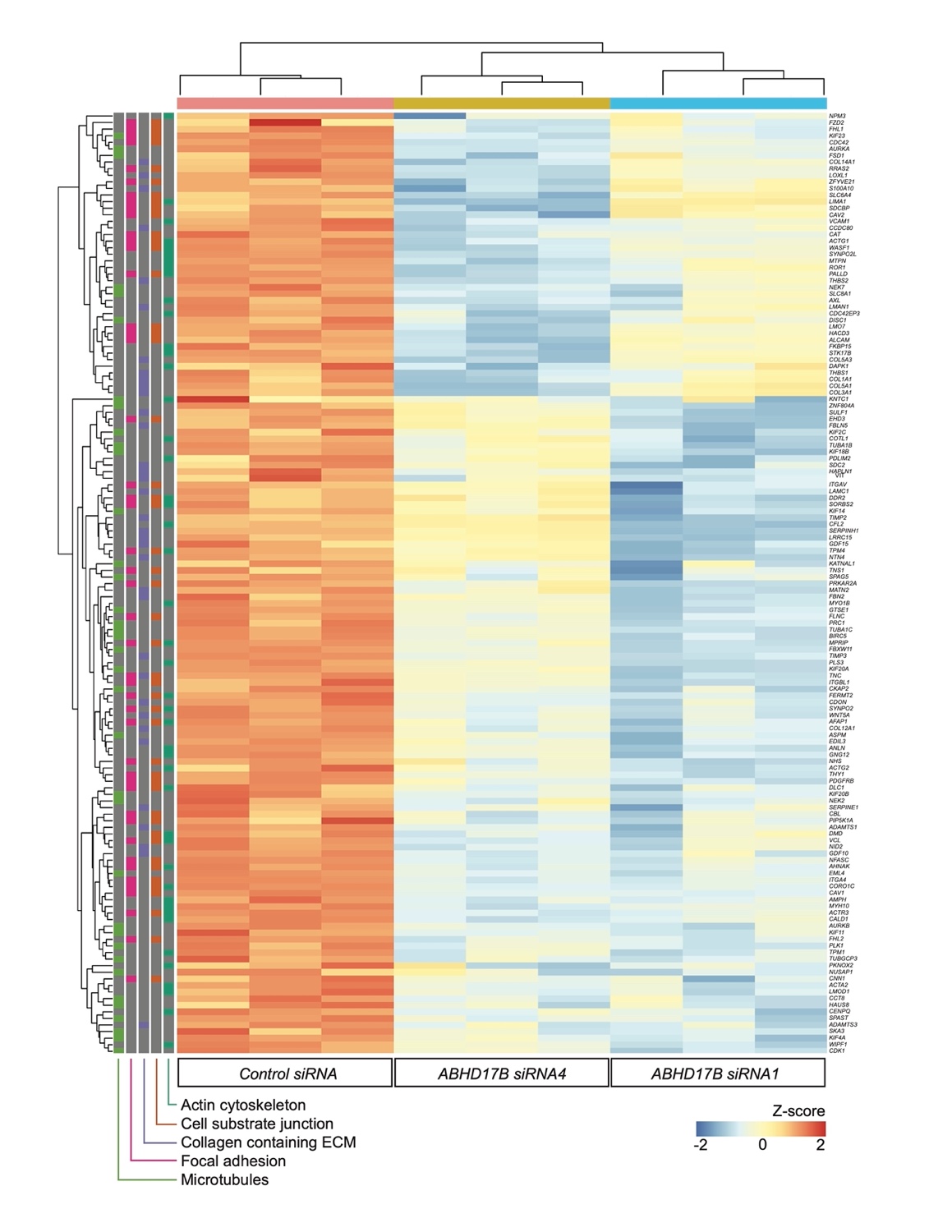

**Supplementary Fig. 12. Genes repressed with depletion of *ABHD17B*.** Heatmap shows expression of the repressed gene set for indicated GO categories for HSCs transfected with control siRNA, siRNA #4 targeting *ABHD17B* and siRNA #1 targeting *ABHD17B.* Gene names are shown on the right. GO categories are indicated by color in the first five gray columns on the left. Expression is centered and scaled by row (gene). The Z-score scale for expression is shown in the lower right.

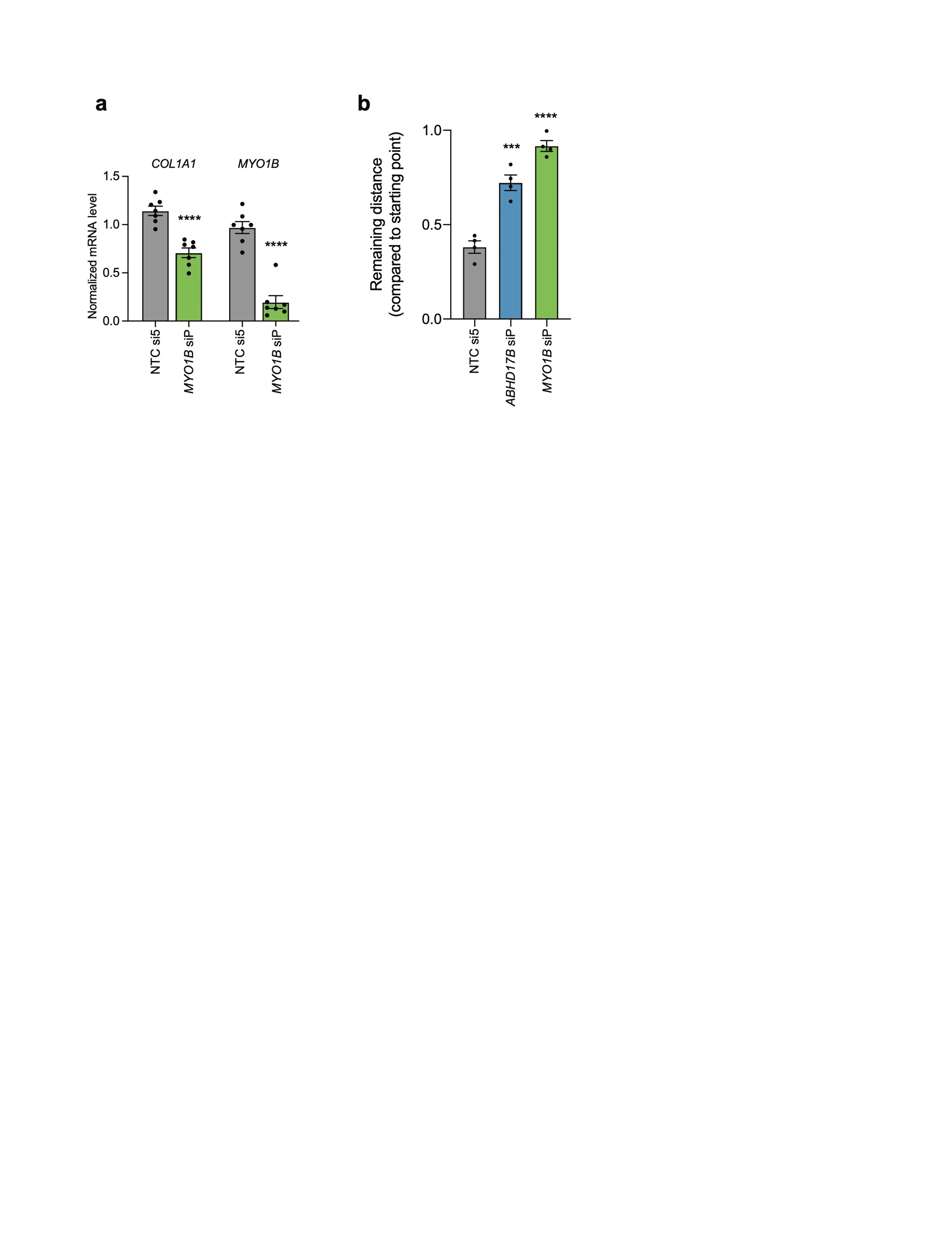

**Supplementary Fig. 13. *MYO1B* depletion reduces *COL1A1* expression and inhibits migration in additional donor lines.** (a) Relative mRNA expression was quantified by qRT-PCR in primary human HSCs (Donor 3) treated witih non targeting siRNAs (NTC si5) and pooled siRNAs targeting *MYO1B*. **** indicates p< 0.0001 (2-tailed unpaired t-test). Data are representative of three independent experiments. (b) Wound healing assay was performed in HSCs (Donor 2) transfected with indicated siRNAs. Normalized wound width was calculated at 88 hr from three individual scratches. *** indicates p < 0.001 and **** indicates p< 0.0001 (one-way ANOVA test).

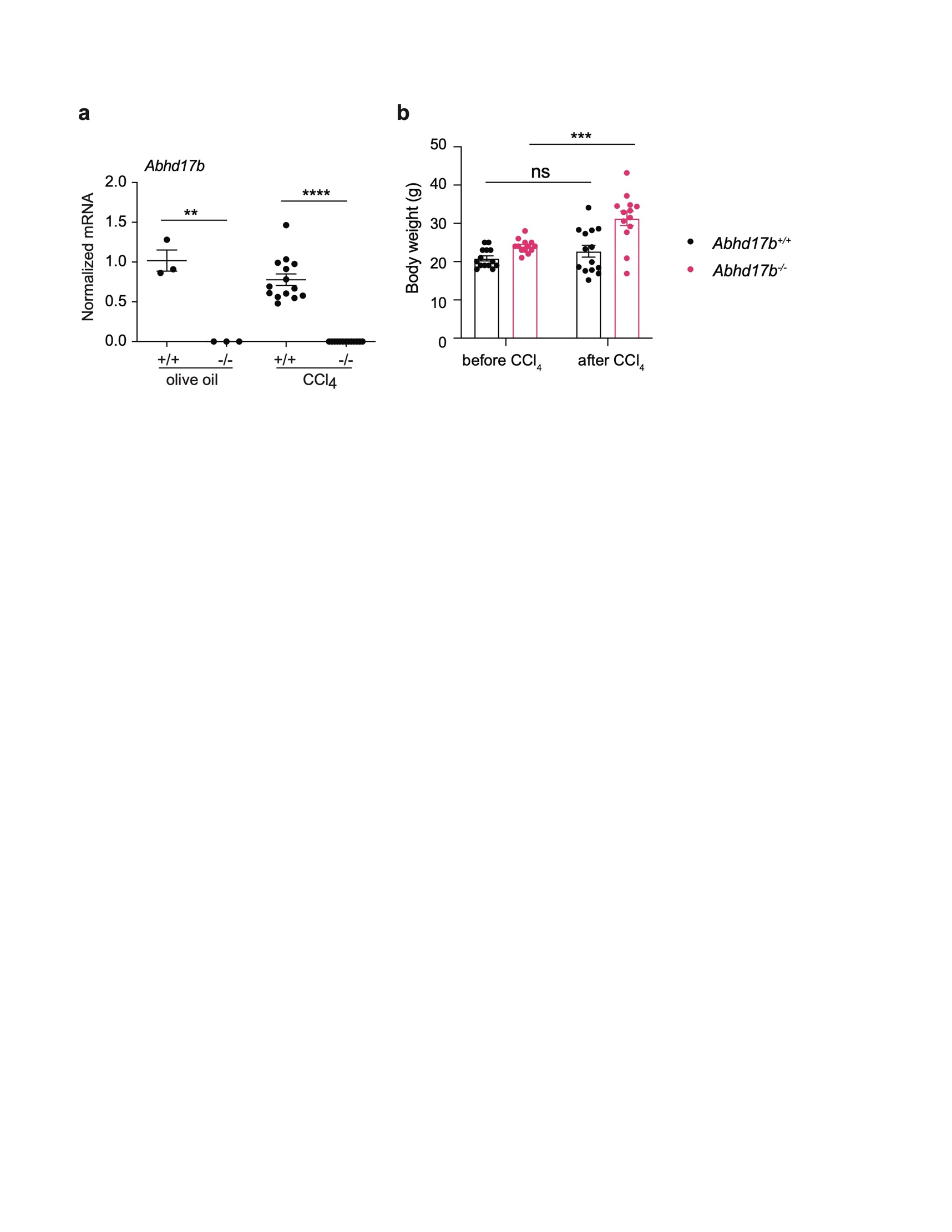

**Supplementary Fig. 14. *Abhd17b* expression in *Abhd17b*-deficient mice and weight changes with CCl_4_ treatment.** (a) Relative *Abhd17b* mRNA expresion was analyzed by qRT-PCR from liver samples. Error bars represent mean ± SEM (n=3, 3, 14, 13), ** indicates p<0.01 and **** indicates p<0.0001 (2-tailed unpaired t-test). (b) Body weight of Abhd17b^+/+^ and Abhd17b^-/-^ mice before or after CCl_4_ treatment. Error bars represent mean ± SEM (n=14, 13), ns indicates not significant, and *** indicates p<0.001 (2-tailed unpaired t-test).

**a**

*ABHD17B*

ATGAATAATCTTTCATTTAGTGAGCTATGTTGCCTCTTCTGCTGTCCACCTTGTCCAGGGAA

GATTGCTTCAAAATTAGCGTTTTTGCCACCTGATCCAACTTACACACTGATGTGTGATGAAA

GCGGAAGCCGTTGGACTTTACATCTGTCTGAACGAGCAGACTGGCAGTATTCTTCTAGAGA

AAAAGATGCTATTGAGTGTTTCATGACTAGAACCAGTAAAGGCAACAGAATTGCTTGTATGT

TTGTACGTTGTTCACCCAATGCGAAATACACTTTACTCTTCTCACATGGAAATGCTGTTGAT

CTTGGTCAAATGAGCAGCTTTTACATAGGACTAGGATCACGGATTAATTGTAATATATTCTC

ATATGATTATTCTGGATATGGTGCCAGTTCCGGGAAACCCACAGAGAAGAACCTCTATGCA

GACATTGAAGCTGCTTGGCTTGCTCTTAGGACAAGATATGGCATTCGCCCTGAAAATGTGA

TTATATATGGCCAAAGTATAGGGACAGTACCGTCTGTGGATCTTGCTGCTCGATATGAGAG

TGCTGCTGTTATTCTTCATTCTCCTCTGACTTCGGGAATGCGAGTTGCCTTTCCaGAcACCA

AGAAGACCTACTGTTTTGATGCATTCCCAAACATTGACAAAATCTCTAAGATAACCTCTCCtG

TtcTAATAATTCATGGGACTGAAGATGAAGTCATTGACTTTTCACATGGCCTCGCATTGTTTG

AACGTTGCCAAAGACCTGTGGAGCCTCTCTGGGTTGAAGGAGCAGGTCACAATGATGTGG

AACTTTATGGACAGTATCTTGAAAGGTTGAAACAGTTTGTGTCACAGGAACTGGTAAATTTG

**b**

*ABHD17B-S170A*

ATGAATAATCTTTCATTTAGTGAGCTATGTTGCCTCTTCTGCTGTCCACCTTGTCCAGGGAA

GATTGCTTCAAAATTAGCGTTTTTGCCACCTGATCCAACTTACACACTGATGTGTGATGAAA

GCGGAAGCCGTTGGACTTTACATCTGTCTGAACGAGCAGACTGGCAGTATTCTTCTAGAGA

AAAAGATGCTATTGAGTGTTTCATGACTAGAACCAGTAAAGGCAACAGAATTGCTTGTATGT

TTGTACGTTGTTCACCCAATGCGAAATACACTTTACTCTTCTCACATGGAAATGCTGTTGAT

CTTGGTCAAATGAGCAGCTTTTACATAGGACTAGGATCACGGATTAATTGTAATATATTCTC

ATATGATTATTCTGGATATGGTGCCAGTTCCGGGAAACCCACAGAGAAGAACCTCTATGCA

GACATTGAAGCTGCTTGGCTTGCTCTTAGGACAAGATATGGCATTCGCCCTGAAAATGTGA

TTATATATGGCCAAGCCATAGGGACAGTACCGTCTGTGGATCTTGCTGCTCGATATGAGAG

TGCTGCTGTTATTCTTCATTCTCCTCTGACTTCGGGAATGCGAGTTGCCTTTCCaGAcACCA

AGAAGACCTACTGTTTTGATGCATTCCCAAACATTGACAAAATCTCTAAGATAACCTCTCCtG

TtcTAATAATTCATGGGACTGAAGATGAAGTCATTGACTTTTCACATGGCCTCGCATTGTTTG

AACGTTGCCAAAGACCTGTGGAGCCTCTCTGGGTTGAAGGAGCAGGTCACAATGATGTGG

AACTTTATGGACAGTATCTTGAAAGGTTGAAACAGTTTGTGTCACAGGAACTGGTAAATTTG

**Supplementary Fig. 15.** **Nucleotide sequence for *ABDH17B* and *ABHD17B-S170A*.** (a) The nucleotide sequence used to encode the wildtype (WT) amino acid sequence for ABHD17B is shown. The position of Ser 170 is indicated in red. Nucleotides in lower case blue text indicate changes from the endogenous nucleotide sequence that do not affect the amino acid sequence. The FLAG peptide was expressed at the C-terminus (not show). (b) The nucleotide sequence was mutated to change Ser 170 to Ala (indicated in red) for ABHD17B-S170A.

References

1. Li, W. *et al.* Nanchangmycin regulates FYN, PTK2, and MAPK1/3 to control the fibrotic activity of human hepatic stellate cells. *Elife* **11**, e74513 (2022).

2. Yang, W. *et al.* Single‐Cell Transcriptomic Analysis Reveals a Hepatic Stellate Cell–Activation Roadmap and Myofibroblast Origin During Liver Fibrosis in Mice. *Hepatology* **74**, 2774–2790 (2021).

3. Patro, R., Duggal, G., Love, M. I., Irizarry, R. A. & Kingsford, C. Salmon provides fast and bias-aware quantification of transcript expression. *Nat Methods* **14**, 417–419 (2017).

4. Soneson, C., Love, M. I. & Robinson, M. D. Differential analyses for RNA-seq: transcript-level estimates improve gene-level inferences. *F1000research* **4**, 1521 (2016).

5. Love, M. I., Huber, W. & Anders, S. Moderated estimation of fold change and dispersion for RNA-seq data with DESeq2. *Genome Biol* **15**, 550 (2014).

6. Wickham, H. *et al.* Welcome to the Tidyverse. *J Open Source Softw* **4**, 1686 (2019).

7. Zhu, A., Ibrahim, J. G. & Love, M. I. Heavy-tailed prior distributions for sequence count data: removing the noise and preserving large differences. *Bioinformatics* **35**, 2084–2092 (2019).

8. Kolde & Heatmaps., Raivo. 2015. pheatmap: P.

9. Yu, G., Wang, L.-G., Han, Y. & He, Q.-Y. clusterProfiler: an R Package for Comparing Biological Themes Among Gene Clusters. *Omics J Integr Biology* **16**, 284–287 (2012).

10. Yu, G. Gene Ontology Semantic Similarity Analysis Using GOSemSim. *Methods Mol Biology Clifton N J* **2117**, 207–215 (2020).

11. Yu, G. *et al.* GOSemSim: an R package for measuring semantic similarity among GO terms and gene products. *Bioinformatics* **26**, 976–978 (2010).

12. Yu, G., Wang, L.-G., Yan, G.-R. & He, Q.-Y. DOSE: an R/Bioconductor package for disease ontology semantic and enrichment analysis. *Bioinformatics* **31**, 608–609 (2015).

13. Ikenaga, N. *et al.* Selective targeting of lysyl oxidase-like 2 (LOXL2) suppresses hepatic fibrosis progression and accelerates its reversal. *Gut* **66**, 1697 (2017).

14. Sojoodi, M. *et al.* Peroxidasin Deficiency Re-programs Macrophages Toward Pro-fibrolysis Function and Promotes Collagen Resolution in Liver. *Cell Mol Gastroenterology Hepatology* **13**, 1483–1509 (2022).

15. Chen, J. Y. *et al.* Tricyclic Antidepressants Promote Ceramide Accumulation to Regulate Collagen Production in Human Hepatic Stellate Cells. *Scientific Reports* **7**, 44867–13 (2017).

16. Yang, J. *et al.* The I-TASSER Suite: protein structure and function prediction. *Nat Methods* **12**, 7–8 (2015).

17. Jumper, J. *et al.* Highly accurate protein structure prediction with AlphaFold. *Nature* **596**, 583–589 (2021).

18. Harder, E. *et al.* OPLS3: A Force Field Providing Broad Coverage of Drug-like Small Molecules and Proteins. *J. Chem. Theory Comput.* **12**, 281–296 (2016).
